# Capturing dynamic fear experiences in naturalistic contexts: An ecologically valid fMRI signature integrating brain activation and connectivity

**DOI:** 10.1101/2023.08.18.553808

**Authors:** Feng Zhou, Ran Zhang, Shuxia Yao, Debo Dong, Pan Feng, Georg Kranz, Tingyong Feng, Benjamin Becker

## Abstract

Enhancing our understanding of how the brain constructs conscious emotional experiences within dynamic real-life contexts necessitates ecologically valid neural models. Here, we present evidence delineating the constraints of current fMRI activation models in capturing naturalistic fear dynamics. To address this challenge, we fuse naturalistic fMRI with predictive modeling techniques to develop an ecologically valid fear signature that integrates activation and connectivity profiles, allowing for accurate prediction of subjective fear experience under highly dynamic close-to-real-life conditions. This signature arises from insights into the crucial role of distributed brain networks and their interactions in emotion modulation, and the potential of network-level information to improve predictions in dynamic contexts. Across a series of investigations, we demonstrate that this signature predicts stable and dynamic fear experiences across naturalistic scenarios with heightened sensitivity and specificity, surpassing traditional activation– and connectivity-based signatures. Notably, the integration of affective connectivity profiles enables accurate real-time predictions of fear fluctuations in naturalistic settings. Additionally, we unearth a distributed yet redundant brain-wide representation of fear experiences. Subjective fear is encoded not only by distributed cortical and subcortical regions but also by their interactions, with no single brain system conveying substantial unique information. Our study establishes a comprehensive and ecologically valid functional brain architecture for subjective fear in dynamic environments and bridges the gap between experimental neuroscience and real-life emotional experience.

## Introduction

Contemporary neuroscience models propose that emotions promote survival in dynamic environments and that the corresponding emotional state is mediated by the interplay between distributed cortical and subcortical brain networks(1–4). Although translational animal models have effectively identified brain regions and neural circuits involved in regulating fear responses in naturalistic contexts(5–9), emerging evidence indicates that the neural mechanisms underlying conscious fear differ from those governing hard-wired behavioral and physiological fear responses(10, 11). This distinction is crucial as animal studies cannot provide insights into the conscious and highly subjective emotional experience that essentially characterizes typical and pathological fear in humans(12–14).

Our understanding of the conscious experience of fear mainly relies on conventional functional Magnetic Resonance Imaging (fMRI) studies in humans. These studies usually employ univariate analysis to localize regional brain activation or connectivity changes elicited by sparse presentations of affective stimuli (e.g., pictures of fearful faces or threatening animals) in laboratory settings. However, the limitations of this localization approach, including moderate effect sizes in brain-outcome associations, have become clear(15–17). To overcome these limitations, fMRI has been combined with multivariate pattern-recognition techniques to establish activation-based brain models for complex subjective experiences including pain(18, 19), negative affect(20, 21) and fear(10, 11). While these models have enabled the development of more precise and comprehensive neural models for emotional experiences(10, 11, 20–25), their capacity for ecologically valid prediction of dynamic emotional experiences in real-world settings remains unexplored. Given the significant differences between laboratory settings and natural environments where fear naturally arises(26), it is imperative to evaluate the sensitivity, specificity, and generalizability of fear-related signatures in predicting dynamic emotional experiences in everyday life, and to establish an ecologically valid neural model for subjective fear in natural environments(2–4, 21, 27). Furthermore, although the network perspective of brain organization has proposed that mental processes arise not from isolated brain regions but from the interplay among multiple areas(3, 28), little is known about the role of brain pathways in encoding subjective experiences.

Utilizing naturalistic stimuli, such as movies, has recently allowed to test experimental brain models under more ecologically valid conditions. This has facilitated the translation of laboratory neuroimaging research into more generalizable brain mechanisms and promoted the exploration of dynamic interactions between brain systems during mental processes(29–32). Movies offer a multisensory experience that can – to a certain extent – simulate the processing of real-life sensory input and embed threat encounters into dynamic and immersive scenarios. This renders movies a more naturalistic and emotionally engaging form of experimental paradigms compared to the conventional presentation of isolated stimuli such as affective pictures. Moreover, evidence suggests that predictive models utilizing network-level information, such as functional connectivity, may better capture attentional and affective changes in dynamic naturalistic environments(33–36), aligning with the importance of dynamic brain system interactions in animal models(1–5, 37).

The present study thus encompasses four main objectives: (1) to test whether previously established fear-related activation-based signatures can predict dynamic fear in naturalistic environments, (2) to develop a sensitive, specific and generalizable fMRI-based signature predictive of stable fear experience in naturalistic contexts while simultaneously capitalizing on whole-brain connectivity and activity (i.e., a synergistic signature), (3) to test whether the synergistic signature can accurately capture fear in dynamic naturalistic environments, and (4) to determine how the subjective experience of fear in naturalistic contexts is represented in brain systems and pathways (Fig. 1).

**Fig. 1.**
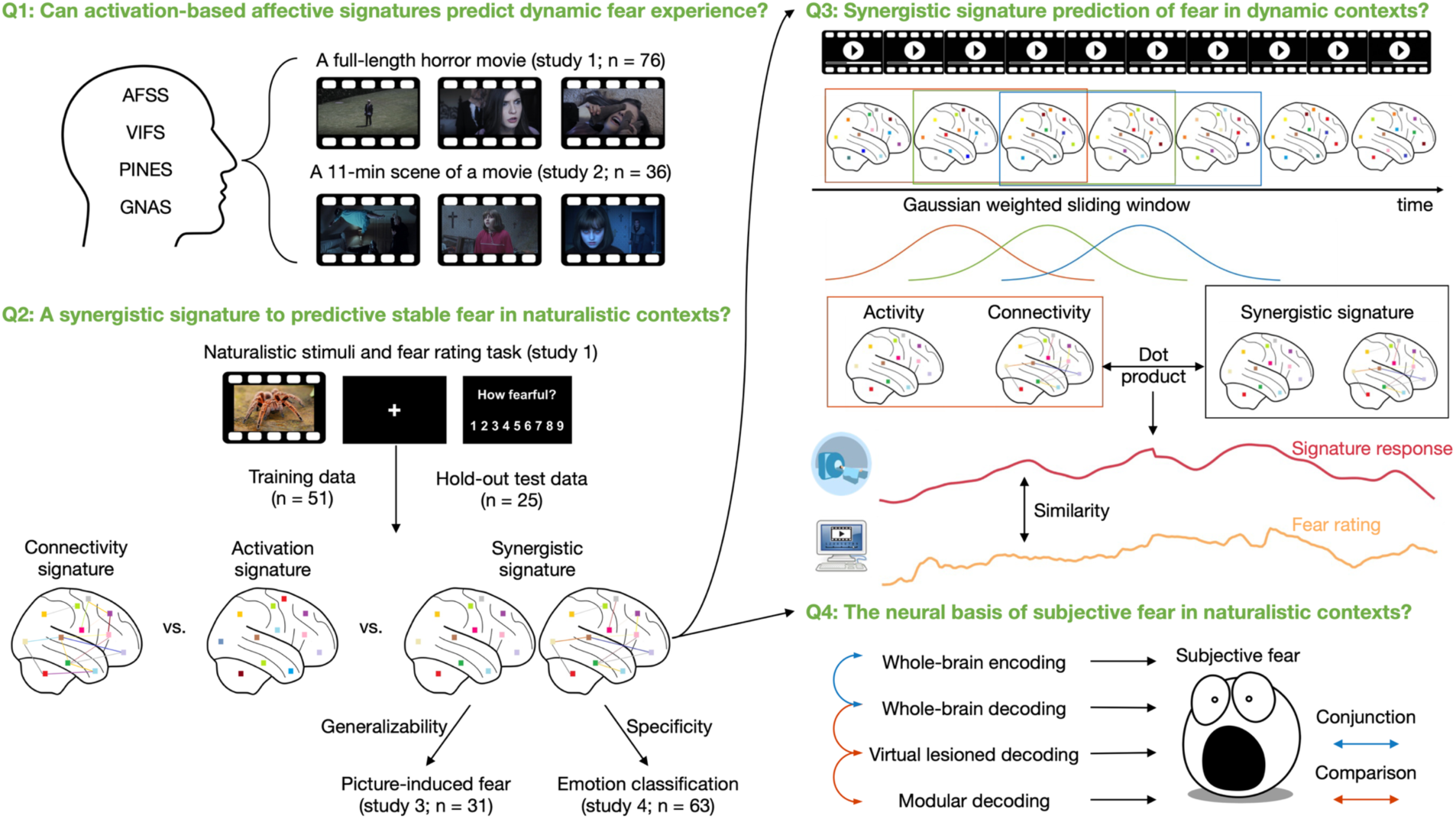
Key research questions, naturalistic paradigms, and synergistic neural decoding of subjective fear experience. We first determined whether the established activation-based fMRI models for fear-related emotions (fear: AFSS and VIFS; general negative affect: PINES and GNAS) could capture dynamic fear under naturalistic conditions (i.e., while watching a full-length horror movie “Don’t look away” and a segment from the horror movie “The Conjuring 2”) (Q1). Next, we combined data from fear induction via immersive movie clips with multivariate predictive modeling to develop a synergistic signature capitalizing on distributed activation and connectivity features to predict fear during a stable emotional experience with higher accuracy than conventional activation and connectivity signatures (Q2). We tested whether the synergistic decoder could track fear variations in the highly dynamic context of watching horror movies (Q3) and further determined which and how brain regions and circuits contribute to the conscious experience of fear in naturalistic contexts (Q4). The examples of the fear-evoking movie clips include pictures only for display purposes and were not included in the original stimulus set. The pictures have been obtained from pixabay.com under the Pixabay License and are free for commercial and noncommercial use across print and digital. AFSS, animal fear schema signature; VIFS, visually induced fear signature; PINES, picture-induced negative emotion signature; GNAS, generalized negative affect signature.

To investigate the predictive capabilities of established activation-based fear signatures (animal fear schema signature, AFSS(11); visually induced fear signature, VIFS(10)) and negative affect signatures (picture-induced negative emotion signature, PINES(20); generalized negative affect signature, GNAS(21)) in capturing dynamic subjective fear experience we conducted two independent fMRI studies. Study 1 (n = 76) involved watching a full-length short horror movie (“Don’t Look Away”; 7 min 28 s), while study 2 (n = 36) focused on a segment from the movie “The Conjuring 2” (11 min 18 s). It’s noteworthy that while the AFSS, VIFS and PINES were originally designed for visual stimuli, the GNAS was developed using a variety of aversive stimuli, including painful heat, painful pressure, aversive images, and aversive sounds, and has previously demonstrated accurate predictions of negative emotions elicited by diverse types of stimuli(21).

In order to enhance the accuracy of predicting both short-term homogeneous and long-term dynamic fear experiences in naturalistic contexts and gain comprehensive insights into the underlying brain regions and pathways of subjective fear, we developed a whole-brain parcel-wise connectivity– and activity-based fear signature (CAFE). This signature was established using a naturalistic fMRI dataset incorporating subjective fear experience reports from study 1 during which participants immersed in 38 video clips of approximately 40 seconds each. We evaluated the predictive performance of the CAFE across studies 1 and 2 and compared it with whole-brain voxel-wise activity and parcel-wise functional connectivity signatures. In study 3 (n = 31), we evaluated the predictive capacity of the CAFE’s activation features, which constitute a small fraction of total features. Furthermore, we examined the generalizability of CAFE and the contributions of arousal and non-specific negative affect in an independent fMRI study (study 4, n = 63) during which subjects rated their arousal while watching 40 naturalistic movie clips (∼30 s) depicting neutral, high arousing negative (fearful, disgusting), or positive events. Finally, through a series of analyses, we elucidated the brain regions and pathways involved in encoding ecologically valid fear experiences in dynamic contexts (Fig. 1).

Overall, the current study allowed us to determine a comprehensive functional brain architecture for subjective fear in naturalistic contexts and provide an ecologically valid brain-based biomarker predictive of subjective fear intensity with high sensitivity and specificity.

## Results

### Previously developed fear-related signatures hardly predict dynamic fear experience during viewing horror movies

We initially tested whether the established activation-based signatures for fear (AFSS(11) and VIFS(10)) and negative affect (PINES(20) and GANS(21)) that were developed on sparsely presented visual(10, 11, 20) or multimodal (visual, somatic and auditory)(21) stimuli could effectively capture dynamic fear experiences during naturalistic movie viewing (Fig. 1).

To this end, we asked subjects in study 1 to watch a short horror movie (7 min 28 s) during fMRI. To approach a naturalistic and ecologically valid dynamic fear experience and associated brain responses no response was required from the subjects during the movie. An additional independent group (n = 30) provided ongoing subjective fear ratings, and after confirming that the trajectory of subjective fear experience induced by the movie is similar across participants (mean ± S.E.M. r = 0.73 ± 0.05), we used the group-average fear rating as a proxy for dynamic fear experience across individuals. Intriguingly, the responses of AFSS, VIFS, PINES, and GNAS showed no significant correlation with (hemodynamic response function) convolved or unconvolved dynamic fear ratings (r ≤0.07, permutation P ≥0.097; r ≤0.07, permutation P ≥0.092, respectively). This observation remained consistent (r ≤ 0.09, permutation P ≥ 0.319) when applying these signatures to the naturalistic data using a tapered sliding window approach(35, 38) (see Fig. 1 for the schematic of this analysis).

We validated these results, in an independent naturalistic fMRI experiment (study 2) during which participants watched a movie segment from another horror movie during fMRI (“The Conjuring 2”, 11 min 18s; n = 36) or provided subjective fear ratings (n = 30), respectively. Similar to prior findings, the trajectory of subjective fear experiences induced by the movie was consistent across participants (r = 0.70 ± 0.05). Correspondingly, the convolved or unconvolved average fear ratings at the group level failed to significantly correlate with the responses of any of the established signatures (r ≤ 0.18, permutation P ≥ 0.056). Taken together, these findings underscore the limitations of neuroaffective signatures solely relying on brain activation magnitude in accurately predicting the nuanced fluctuations in emotional experiences within complex real-world contexts.

### A synergistic brain connectivity– and activity-based signature for subjective fear (CAFE) in naturalistic contexts

Our study aimed to develop an innovative fMRI-based decoder that can accurately predict both short-term stable and long-term dynamic fear experiences. Specifically, we focused on predicting responses to short movie clips that reliably elicit consistent subjective fear, as well as responses to longer films that induce fluctuating fear encounters. By leveraging our understanding of the involvement of distributed brain networks and their interplay in emotional modulation(1–4) and the potential of network-level information (functional connectivity) to enhance predictions in dynamic environments(33–36), we hypothesized that combining whole-brain activity and functional connectivity profiles would improve prediction accuracy in both stable and dynamic fear experiences within naturalistic contexts.

To this end, n = 76 participants in study 1 were shown a total of 38 immersive movie clips lasting 30-50 s and reported their level of subjective fear on a scale ranging from 1 to 9 during fMRI. Importantly, the movie clips effectively induced a wide range of subjective fear levels, while maintaining consistent emotional intensity throughout each video. This allowed us to capture reliable mean activation and functional connectivity for each fear intensity level. Participants were split into training (n = 51) and test (n = 25) datasets. Subsequently, we employed the linear support vector regression (SVR) algorithm and developed a synergistic brain connectivity– and activity-based signature for subjective fear (i.e., the CAFE), which incorporated both parcel-mean activation and functional connectivity between all pairs of regions as its constituent features, using the training data alone. Considering evidence from prior investigations suggesting that the human neocortex can be segregated into 300-400 functionally separable areas(39, 40), we developed the CAFE using a parcellation scheme comprising a total of 463 parcels, encompassing cerebral, subcortical, and cerebellar regions (see Methods for details).

To assess the efficacy of the CAFE we employed 10 repeats of 10-fold cross-validation on the training data, followed by out-of-sample predictions on the test data. As shown in Fig. 2a, the CAFE accurately predicted the stable subjective fear experience elicited by movie clips, both in the training dataset (cross-validated prediction-outcome correlation r = 0.64, R^2^ = 0.40, both permutation P < 1 × 10^-4^) and the test dataset (r = 0.67, R^2^ = 0.43, both permutation P < 1 × 10^-4^). The synergistic fear model, CAFE, surpassed the conventional whole-brain voxel-wise activation-based signature (training data: r = 0.60, R^2^ = 0.33; test data: r = 0.62, R^2^ = 0.36) to a varying degree, with the differences of R^2^ achieving (marginal) significance (training data: permutation P = 0.006; test data: permutation P = 0.085). Importantly, the CAFE statistically outperformed the connectivity-based signature (training data: r = 0.58, R^2^ = 0.33; test data: r = 0.60, R^2^ = 0.34) in both training and test datasets (all permutation P ≤ 0.010). We further compared the prediction performance of the CAFE with those of previously established activation-based signatures (AFSS, VIFS, PINES, and GNAS). The CAFE was developed on the training data and to facilitate a fair comparison we therefore focused on the hold-out test dataset (for predictions on the whole sample see Supplementary Fig. 1). We found that the CAFE exhibited superior performance compared to these activation-based signatures (all permutation P < 1 × 10^-4^; see Supplementary Results for details). Notably, the prediction performance of the synergistic signature remained robust when we randomly split the training and test data 100 times, and when we applied a different parcellation method (see Supplementary Results for details).

**Fig. 2.**
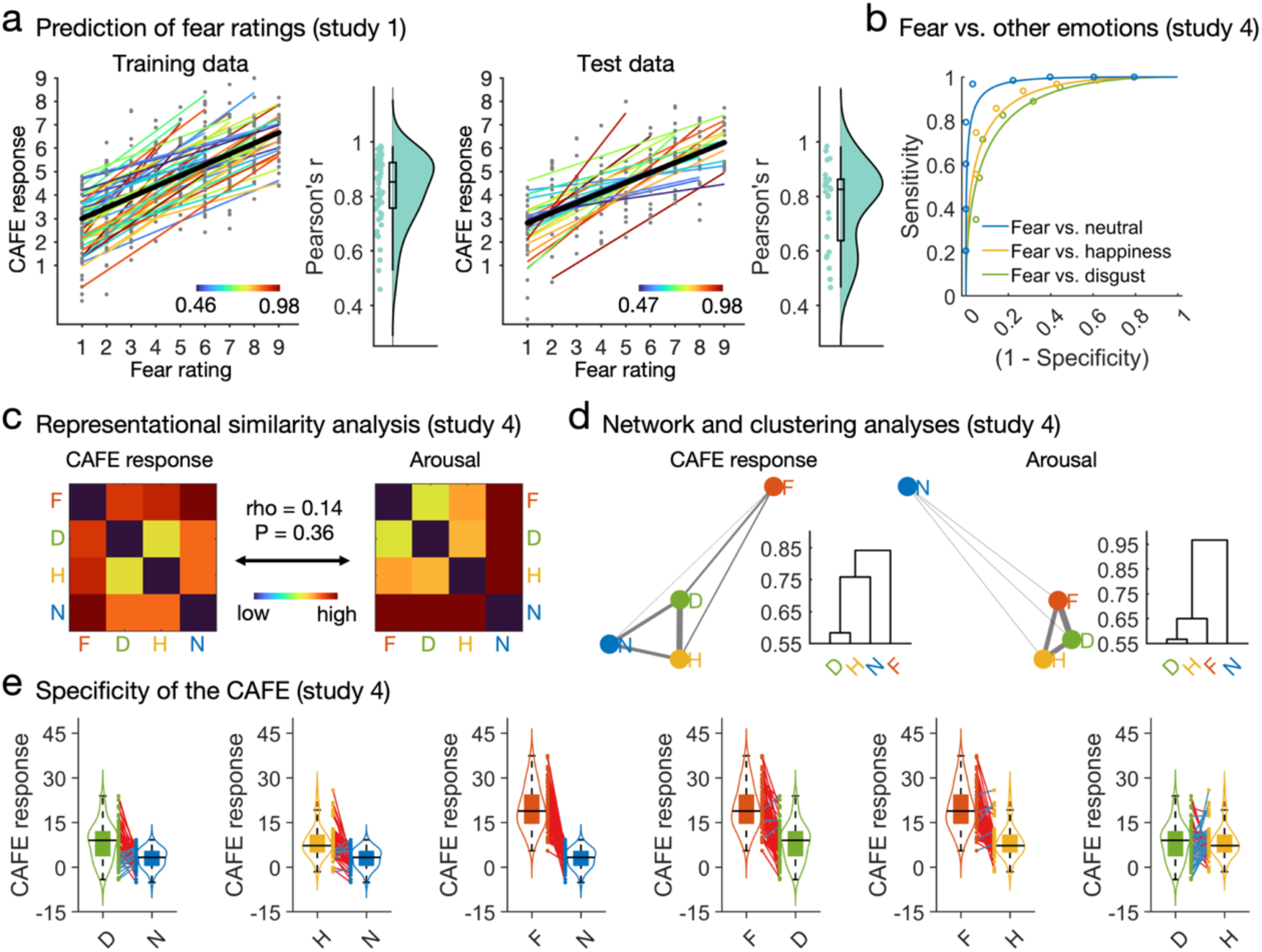
Sensitivity and specificity of the CAFE. a, the CAFE accurately predicts subjective fear experience induced by movie clips (overall prediction-outcome r ≥ 0.64, R^2^ ≥ 0.40, permutation P < 1×10^-4^). Each colored line represents prediction within each individual participant. Black line indicates the overall (i.e., within and between participants) prediction. Raincloud plots show the distribution of within-participant predictions. b, the CAFE accurately distinguishes fear from disgust, happiness and neutral in naturalistic contexts (study 4; accuracies > 86.77%, P < 1.05 × 10^-5^, d > 1.76). Representational (c) and network and clustering (d) analyses show that the fear predictions by the CAFE are distinct from those based on arousal ratings. Correlation between the two matrices were performed by the Mantel test, and P value was obtained with 10,000 permutations. e, the CAFE captures a small extent of arousal, but not negative valence, information. High arousing disgust– and happiness-inducing movie clips exhibit stronger CAFE responses as compared to low arousing neutral clips (accuracy = 75.83%, P = 1 × 10^-8^, d = 1.04; accuracy = 75.83%, P = 1 × 10^-8^, d = 1.24, respectively). However, the CAFE predicts fear vs. neutral (accuracy = 97.50%, P < 1 × 10^-20^, d = 2.97) considerably more accurate as reflected by effect sizes 2.39-2.85 times larger than those for predicting disgust or happiness vs. neutral. Moreover, the CAFE can accurately classify fear from arousal-matched disgust (accuracy = 84.17%, P = 1 × 10^-14^, d = 1.68) and (close-to) matched happiness (accuracy = 88.33%, P = 1 × 10^-20^, d = 1.79). Each colored line between dots represents each individual participant’s paired data (red line indicates correct classification whereas blue line indicates incorrect classification based on the forced two-choice test). F, fear; D, disgust; H, happiness; N, neutral.

Interestingly, despite comprising a small fraction (∼0.4%) of total features, the incorporation of activation features bolstered the CAFE’s predictive capability in comparison to connectivity-based signature. However, the extent to which the activation pattern of the CAFE can predict subjective fear experiences remains uncertain. To address this issue, we applied the activation pattern of the CAFE to an independent fMRI dataset consisting of n = 31 participants who underwent fMRI scanning while viewing animal and object pictures. Before the fMRI procedure, subjective fear ratings were obtained for each animal category without presenting any stimuli(11). Our results revealed a significant predictive relationship between the response of CAFE’s activation pattern and the subjective fear ratings for animal categories (n = 163 brain images, r = 0.35, P = 6 × 10^-6^; Supplementary Fig. 2). These findings suggest that the activity features play a significant role in the synergistic fear model, contributing to the superior performance of the CAFE model when compared to the connectivity-based model.

### The CAFE generalizes across contexts and depends to a small extent on arousal, but not non-specific negative affect

One of the main challenges in predictive modeling of emotional signatures is determining whether the signature specifically captures discrete emotional states such as fear, as opposed to general and nonspecific emotional processes like salience, valence, or arousal, which are inherently related to the emotional process of interest(27, 41, 42). To address this critical issue and examine the generalizability of the CAFE, we applied the CAFE to fMRI data collected in study 4 showing short movie clips to elicit a range of positive and negative arousing and non-arousing experiences.

To assess the influence of arousal on the classification performance of the CAFE, we conducted a series of analyses. We employed exploratory representational similarity analysis, network analysis, and hierarchical clustering(43) to compare classification accuracies between pairs of emotions based on the CAFE signature response and arousal rating separately. Our initial findings demonstrated that the CAFE model exhibited a significant performance to accurately differentiate fear from other emotions (accuracies ≥ 84.17%, P < 1 × 10^-14^; Fig. 2b). Furthermore, as evidenced in Fig. 2c and 2d, the predictions derived from the CAFE model and those rooted in arousal-based assessments manifested discernible patterns. This suggests that the predictive capability of the CAFE model is not exclusively reliant on arousal. Nonetheless, it is noteworthy that the CAFE responses for disgust and happiness surpassed those for neutral stimuli, implying a certain degree of association between the CAFE model and arousal.

To further quantify the extent to which arousal contributed to the CAFE prediction we matched arousal ratings across emotional categories by excluding data from non-neutral movie clips that induced only low-to-medium levels of subjective arousal. This resulted in a dataset with comparable levels of high arousal induction for fear and disgust, although fear clips still induced slightly higher arousal than happiness clips (see Methods for details). As shown in Fig. 2e, heightened CAFE responses were observed for disgust and happiness compared to neutral (75.83%, P = 1 × 10-8, d = 1.04 and 75.83%, P = 1 × 10-8, d = 1.24, respectively). However, contrasting these accuracies with fear vs. neutral predictions (accuracy = 97.50%, P < 1 × 10^-20^, d = 2.97) highlighted the CAFE’s superior prediction for fear, with effect sizes (Cohen’s d) 2.39-2.85 times larger than predicting disgust or happiness. Crucially, the CAFE effectively differentiated fear from arousal-matched disgust (accuracy = 84.17%, P = 1 × 10^-14^, d = 1.68) and near-matched happiness (accuracy = 88.33%, P = 1 × 10^-20^, d = 1.79), effect sizes 1.44-1.61 times larger than predicting disgust and happiness vs. neutral. Notably, findings remained consistent when analyzing the complete dataset with all video clips. These results suggest that while the CAFE captures arousal information inherent to fear, arousal’s impact on prediction performance remains modest.

To explore the dependence of the CAFE on non-specific negative affect, we compared its signature response during non-fear negative (disgust) and positive (happiness) emotional experiences. We hypothesized that if the CAFE captured non-specific negative valence, it would exhibit a higher signature response for disgust compared to happiness. However, the CAFE failed to classify disgust from happiness accurately in either the full dataset (accuracy = 57.94%, P = 0.090, d = 0.13) or the high arousal dataset (accuracy = 58.33%, P = 0.082, d = –0.02). Furthermore, direct comparisons of the signature responses indicated no significant differences between disgust and happiness (P = 0.475, BF_01_ = 5.66 and P = 0.909, BF_01_ = 7.20, respectively; paired t-test).

Additionally, we compared the sensitivity and specificity of the CAFE with connectivity– and activation-based signatures developed in Study 1, as well as established fear-related signatures. Our results demonstrated that the CAFE exhibited significantly higher sensitivity and specificity (see Table 1 for details). For example, while the connectivity-based signature showed comparable accuracy in predicting dynamic fear compared to the CAFE model (see below for details), it had lower prediction rates for distinguishing stable fear from neutral conditions when compared to the CAFE (80.00% vs. 97.50%). Moreover, the connectivity-based signature captured non-specific valence, with significantly stronger activation for disgust clips compared to happiness clips (paired t-test P = 0.005, BF_10_ = 6.79 and P = 0.003, BF_10_ = 10.90 in full and high arousal datasets, respectively). In summary, our findings suggest that the CAFE exhibits a high level of specificity compared to other fear-related signatures, effectively capturing a greater degree of fear-specific information while moderately considering arousal-related factors, without capturing non-specific negative affect.

**Table 1.**
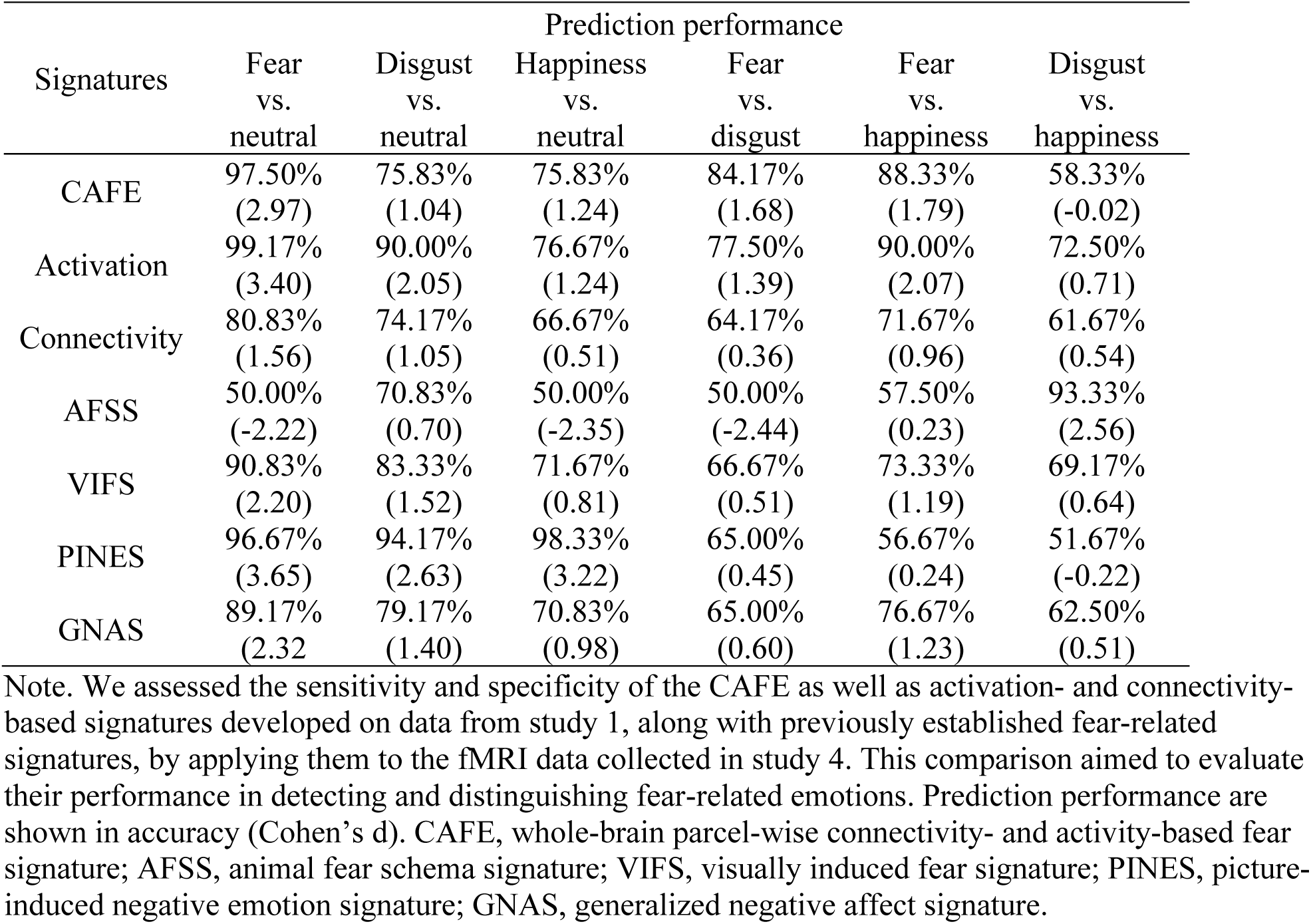
Comparison of prediction performance between the CAFE and signatures developed on study 1 data, as well as previously established fear-related signatures.

| Signatures | Prediction performance |  |  |  |  |  |
| --- | --- | --- | --- | --- | --- | --- |
|  | Fear<br>vs.<br>neutral | Disgust<br>vs.<br>neutral | Happiness<br>vs.<br>neutral | Fear<br>vs.<br>disgust | Fear<br>vs.<br>happiness | Disgust<br>vs.<br>happiness |
| CAFE | 97.50%<br>(2.97) | 75.83%<br>(1.04) | 75.83%<br>(1.24) | 84.17%<br>(1.68) | 88.33%<br>(1.79) | 58.33%<br>(-0.02) |
| Activation | 99.17%<br>(3.40) | 90.00%<br>(2.05) | 76.67%<br>(1.24) | 77.50%<br>(1.39) | 90.00%<br>(2.07) | 72.50%<br>(0.71) |
| Connectivity | 80.83%<br>(1.56) | 74.17%<br>(1.05) | 66.67%<br>(0.51) | 64.17%<br>(0.36) | 71.67%<br>(0.96) | 61.67%<br>(0.54) |
| AFSS | 50.00%<br>(-2.22) | 70.83%<br>(0.70) | 50.00%<br>(-2.35) | 50.00%<br>(-2.44) | 57.50%<br>(0.23) | 93.33%<br>(2.56) |
| VIFS | 90.83%<br>(2.20) | 83.33%<br>(1.52) | 71.67%<br>(0.81) | 66.67%<br>(0.51) | 73.33%<br>(1.19) | 69.17%<br>(0.64) |
| PINES | 96.67%<br>(3.65) | 94.17%<br>(2.63) | 98.33%<br>(3.22) | 65.00%<br>(0.45) | 56.67%<br>(0.24) | 51.67%<br>(-0.22) |
| GNAS | 89.17%<br>(2.32) | 79.17%<br>(1.40) | 70.83%<br>(0.98) | 65.00%<br>(0.60) | 76.67%<br>(1.23) | 62.50%<br>(0.51) |

Note. We assessed the sensitivity and specificity of the CAFE as well as activation- and connectivity-based signatures developed on data from study 1, along with previously established fear-related signatures, by applying them to the fMRI data collected in study 4. This comparison aimed to evaluate their performance in detecting and distinguishing fear-related emotions. Prediction performance are shown in accuracy (Cohen's d). CAFE, whole-brain parcel-wise connectivity- and activity-based fear signature; AFSS, animal fear schema signature; VIFS, visually induced fear signature; PINES, picture-induced negative emotion signature; GNAS, generalized negative affect signature.

### The synergistic signature predicts dynamic subjective fear in naturalistic contexts

In everyday life fear evolves in interaction with dynamic environmental changes and the translational potential of an ecologically valid neural signature critically depends on its capacity to track these variations. We thus explored the potential of the CAFE to capture moment-to-moment variations in subjective fear experiences during the full-length horror movies acquired in studies 1 and 2. Using a tapered sliding window approach, we applied CAFE to the naturalistic movie data collected in study 1, followed by prediction evaluations on both training dataset (cross-validated) and hold-out test dataset. Leveraging the movie’s consistent impact on subjective fear experiences across participants (Fig. 3a), we employed group-average fear ratings as proxies for dynamic fear variations across individuals. We then quantified prediction accuracy by calculating average Fisher’s z-transformed Pearson’s correlations between individually predicted fear dynamics and observed group-level fear ratings(35). As depicted in Figure 3b, the CAFE model accurately predicted subjective fear experience above chance level in both the training dataset (r = 0.37 ± 0.05, permutation P = 2 × 10^-4^) and test dataset (r = 0.44 ± 0.06, permutation P = 4 × 10^-4^). Notably, a robust positive correlation was discovered between the average neural response spanning the duration of the movie and the group-average fear ratings (r = 0.80, permutation P = 0.004 and r = 0.85, permutation P = 0.002, for training and test dataset respectively). Comparatively, the predictive performance of the CAFE model slightly surpassed the connectivity-based model (training data: within-participant prediction-outcome r = 0.34 ± 0.05, permutation P = 0.002, group-average prediction-outcome r = 0.78, permutation P = 0.014; test dataset: within-participant prediction-outcome r = 0.42 ± 0.06, permutation P = 0.001, group-average prediction-outcome r = 0.85, permutation P = 0.006). Significantly, the CAFE model exhibited substantially improved predictive capabilities compared to the activation-based model (training data: within-participant prediction-outcome r = 0.12 ± 0.02, permutation P = 0.061, group-level prediction-outcome r = 0.26, permutation P = 0.0564; test dataset: within-participant prediction-outcome r = 0.20 ± 0.04, permutation P = 0.015, group-average prediction-outcome r = 0.36, permutation P = 0.023).

**Fig. 3.**
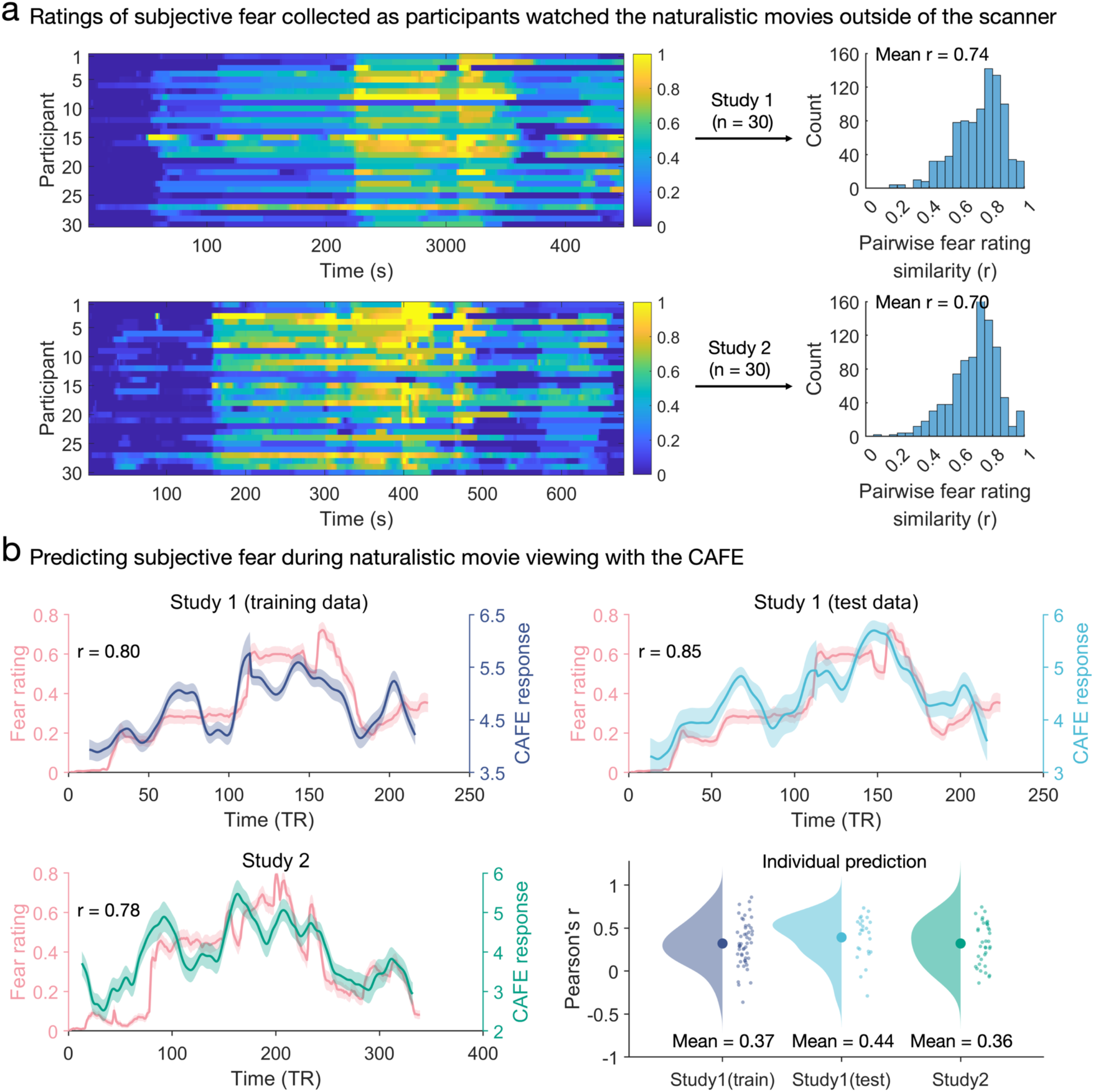
The CAFE predicts group-average dynamic states of subjective fear experience. a, continuous real-time ratings of subjective fearing were collected while participants watched two horror movies (“Don’t look away” and “The Conjuring 2”) in an independent behavioral experiment. The left panel depicts ratings across time for each participant and the right panel depicts histograms of pairwise participants’ response similarities. Mean r values were computed by averaging Fisher’s z – transformed Pearson’s correlation coefficients and transforming the mean Fisher’s z value back to r. b, testing whether the CAFE can accurately predict the dynamic moment-to-moment changes in subjective fear during naturalistic movie viewing on the group level revealed significant associations in both study 1 (training data: r = 0.80, permutation P = 0.004; test data: r = 0.85, permutation P = 0.002) and study 2 (r = 0.78, permutation P = 0.015). Lines represent mean values and shaded areas represent the standard error of mean. Moreover, the lower-right panel shows that the CAFE exhibits significant within-participant predictions in both study 1 (r = 0.37 ± 0.05, permutation P = 2 × 10^-4^ and r = 0.44 ± 0.06, permutation P = 4 × 10^-4^ for training and test datasets, respectively) and study 2 (r = 0.36 ± 0.05, permutation P = 0.003).

These findings were successfully replicated in study 2, further corroborating our initial observations. Specifically, the CAFE predictions exhibited significant correlations with the group-average fear ratings (r = 0.36 ± 0.05, permutation P = 0.003). Additionally, we found a strong positive correlation between the group-average signature response and the group-average fear rating throughout the movie (r = 0.78, permutation P = 0.015). Consistent with our previous results, the prediction performances of the CAFE model were comparable to those of the connectivity-based model (within-participant prediction-outcome r = 0.36 ± 0.05, permutation P = 0.005; group-average prediction-outcome r = 0.77, permutation P = 0.019). Moreover, the CAFE model outperformed the activation-based model substantially (within-participant prediction-outcome r = 0.13 ± 0.03, permutation P = 0.077; group-average prediction-outcome r = 0.27, permutation P = 0.088).

Importantly, the prediction of the fear experience with the CAFE remained robust across different sliding window lengths (Supplementary Fig. 3). In contrast, activation-based fear signature developed using naturalistic stimuli demonstrated better prediction of dynamic fear experiences compared to those developed using static pictures. However, it still exhibited limitations in capturing moment-to-moment changes in subjective fear, thus highlighting the restricted efficacy of neural signatures that solely rely on activation features in capturing the dynamic nature of emotional experiences. Additionally, predicting fear dynamics using the connectivity and activity patterns of the CAFE separately further confirmed that connectivity, but not activity, profiles considerably contributing to the predictive accuracy. Our results emphasize the significance of integrating affective connectivity profiles to enable precise and real-time predictions of fear fluctuations within naturalistic settings.

### Fear experience in natural contexts is represented in distributed brain regions and pathways

In our previous study(10), we identified brain regions associated with and predictive of subjective fear but did not account for dynamic interactions between the systems. Analyzing consistent predictive weights in the synergistic model and examining the encoding map(44) (Supplementary Fig. 4) allowed us to gain insights into specific brain regions and pathways contributing to subjective fear, with or without considering the effects of other regions and circuits, respectively. To identify key brain features underlying fear experience, we focused on features significant in both predictive weights and encoding map (i.e., their conjunction after thresholding at FDR P < 0.05)(10, 21, 44).

Consistent with our prior findings(10), we observed positive weights and associations in distributed cortical and subcortical regions involved in fear reactivity and emotional awareness(14), including the (hypo)thalamus, periaqueductal gray (PAG), brainstem, anterior insula, and midcingulate cortex (MCC). Conversely, regions commonly associated with implicit emotion regulation, interoception, and somatomotor processing(37, 45), such as the ventromedial prefrontal cortex (vmPFC), anterior lateral prefrontal cortex (alPFC), middle temporal gyrus, inferior parietal lobule (IPL), posterior insula/operculum, and somatomotor cortex, exhibited negative weights and associations (Fig. 4a). Moreover, our findings revealed significant connections in both the weights and structure coefficients among all brain parcels, suggesting the involvement of distributed neural pathways in fear experience. Increased functional connectivity with cerebellar and visual cortices, including the PAG and brainstem, primarily predicts heightened fear experience, indicating the involvement of circuits related to threat detection and initial threat reactivity(14). Conversely, connectivity with somatomotor regions strongly predicts decreased subjective fear, potentially reflecting processes involving somatic representation, affective experience, or emotion regulation. At the node level, the MCC displays the highest positive weighted degree, implying that connectivity with this region is indicative of heightened fear. Conversely, connectivity with the dorsal posterior insula (dpINS) predicted lower subjective fear (Fig. 4b).

**Fig. 4.**
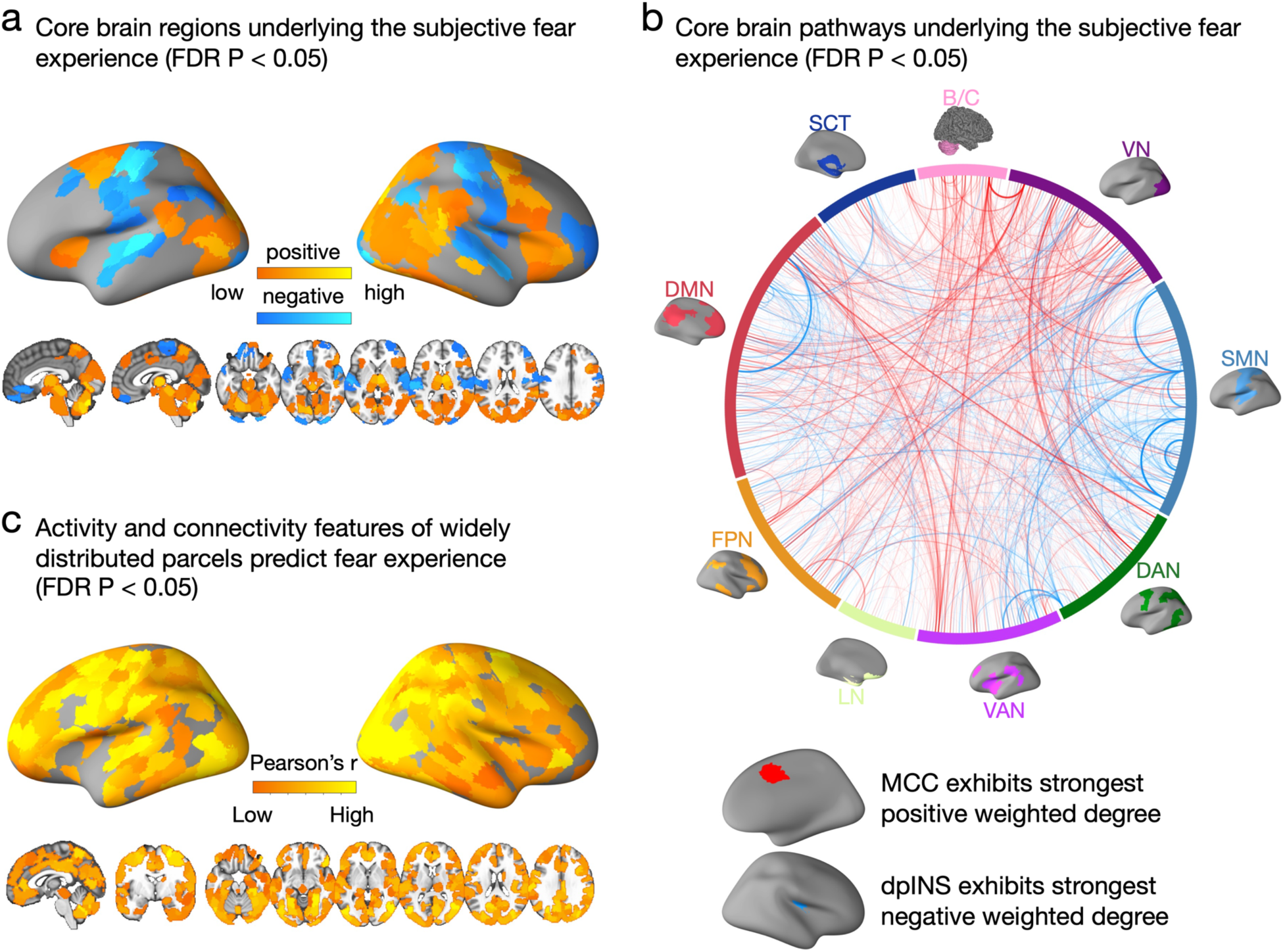
Core brain regions and pathways underlying subjective fear experience in naturalistic contexts. a, core brain regions for the subjective fear experience; defined as the conjunction (FDR P < 0.05) of the synergistic signature weight map and the model encoding map. b, circular plot visualization of core pathways for the subjective fear experience; defined as the conjunction (FDR P < 0.05) of the synergistic signature weight map and the model encoding map. We group brain regions into 9 networks for display purpose. VN, visual network; SMN, somatomotor network; DAN, dorsal attention network; VAN, ventral attention network; LN, limbic network; FPN, frontoparietal network; DMN, default mode network; SCT, subcortical network; B/C, brainstem, and cerebellum; MCC, middle cingulate cortex; dpINS, dorsal posterior insula. c, brain regions which can significantly predict subjective fear ratings in both training (cross-validated) and test datasets by regional activity and connectivity with other regions (FDR P < 0.05).

To elucidate the impact of the synergistic approach on the identification of core brain features, we compared the core connectivity features of the CAFE and the connectivity (only) signature. We focused on this comparison because these signatures share over 99% common features, which could provide significant insights into how the inclusion of different brain features (activation), even in a limited number, can impact the identification of core brain pathways underlying the subjective fear experience. We found that while the unthresholded weight map and the encoding map of the CAFE pathways were highly correlated with those of the connectivity signature (r > 0.80), the synergistic CAFE model is more conservative in identifying core pathways (thresholded) underlying the subjective fear experience. That is, although most core pathways (over 91%) identified in the CAFE remained significant in the connectivity signature the activity features could partially “suppress” the contribution of some pathways in the synergistic model (i.e., only 67% of those core pathways identified in the connectivity model retained their significance in the synergistic model). This “suppression” is particularly noteworthy, considering that the activation features represented only 0.43% of the entire features of the CAFE, and it might reflect the complex interactions between regional activation and cross-regional communication underlying the subjective fear experience (see Supplementary Fig. 5 for details).

To better understand the neurobiological foundations of subjective fear, we conducted further analysis to determine which brain region could predict fear experience based on its activation and connectivity with all other regions. Rather than using statistical inference to test hypotheses about the weights in the comprehensive predictive model, this analysis summarized the evidence using local multivariate statistics (i.e., prediction performance). This approach, similar to traditional univariate brain mapping analysis, allows for interpretability as each region is independently subjected to the same analysis(46). As shown in Fig. 4c, widely distributed cortical and subcortical regions (321 out of 463 parcels), including the vmPFC, MCC, PAG, insula and amygdala, could predict subjective fear in both training (cross-validated) and test data (i.e., their conjunction; each thresholded at FDR P < 0.05 based on permutation tests), further supporting the distributed nature of the neural basis underlying subjective fear.

### Subjective fear experience is encoded and represented in a redundant fashion

To assess the necessity and uniqueness of each brain system in subjective fear, we partitioned the whole-brain into nine large-scale networks, encompassing cortical, subcortical, and cerebellar regions. Subsequently, we randomly selected parcels ranging from 40 to 400 within these networks and utilized their connectivity and activity features to train new predictive models for subjective fear. Fig. 5a displays an improvement in prediction performance as the number of selected parcels increased, particularly when including parcels from a greater number of networks. Statistical analysis using linear regression models confirmed these observations. The number of selected parcels, the number of networks from which they were drawn, and their interaction significantly influenced prediction performances (all P value < 4.76 × 10^-6^, partial η^2^ > 0.53). Remarkably, the prediction performance remained stable when using half or more of the parcels to train the models.

**Fig. 5.**
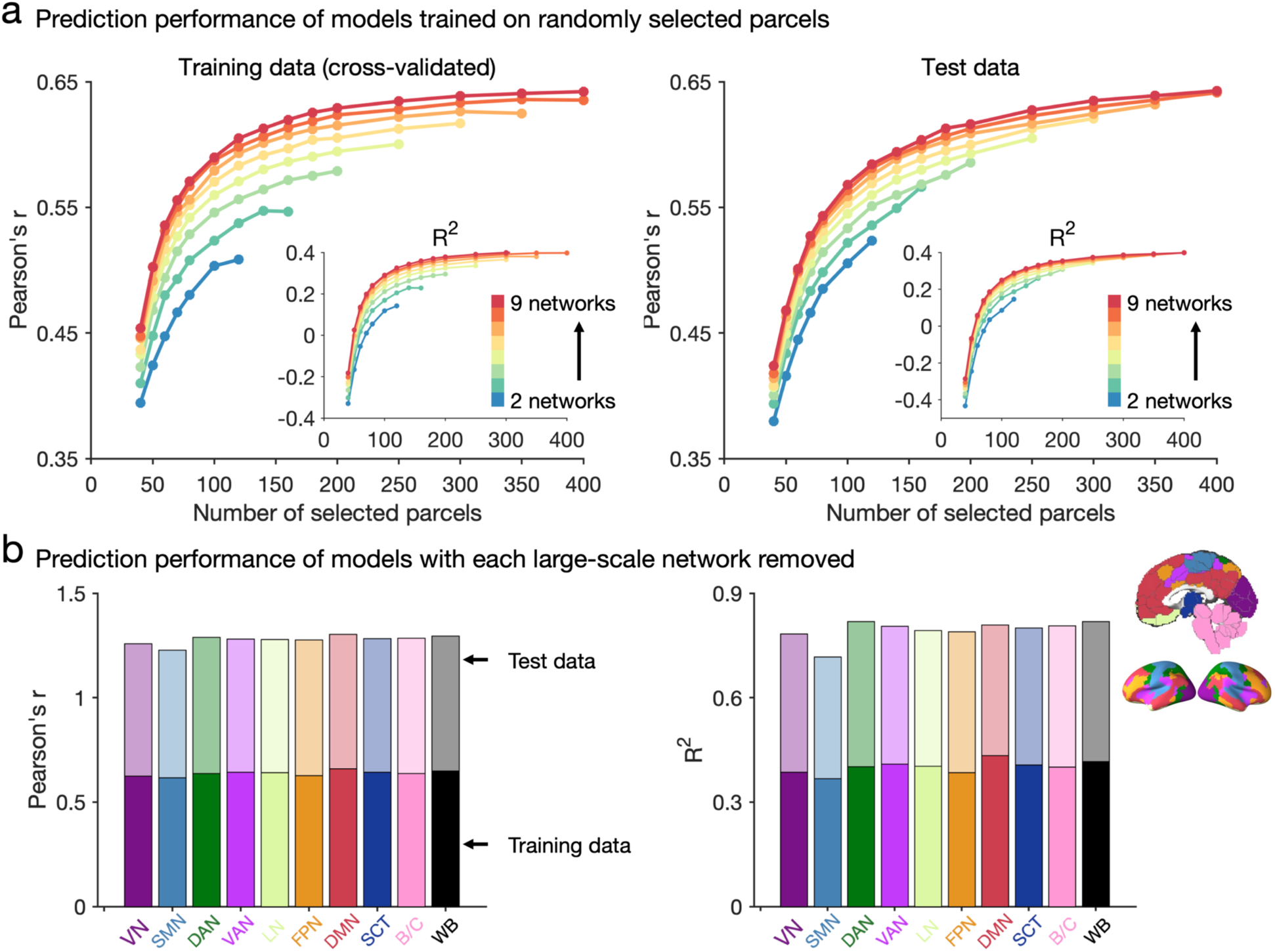
Subjective fear experience is encoded and represented in a redundant fashion. a, predictions based on randomly selected regions show that model performance increases as a function of the numbers of selected parcels and for a given number of selected parcels model performance increases as a function of parcels spanning more networks. Dot indicates the mean prediction performance across 1,000 random samples. However, the prediction performance does not strongly change when only half or more parcels were used to train the models. b, “virtual lesion” analysis illustrates that no single subnetwork is necessary for accurately predicting the intensity of subjective fear. VN, visual network; SMN, somatomotor network; DAN, dorsal attention network; VAN, ventral attention network; LN, limbic network; FPN, frontoparietal network; DMN, default mode network; SCT, subcortical network; B/C, brainstem, and cerebellum; WB, whole brain.

Based on these findings, we hypothesized that subjective fear experience might be redundantly encoded, with each network providing limited unique information. To investigate this, we conducted a comprehensive analysis. Firstly, we employed a “virtual lesion” analysis(20) and demonstrated that removing individual networks had minimal impact on model performance (training data: r = [0.62, 0.66], R^2^ = [0.37, 0.43]; test data: r = [0.61, 0.65], R^2^ = [0.35, 0.42]) (Fig. 5b). In this analysis we re-trained the model when each network was removed. Moreover, our finding was evident even when using a partial pattern of the CAFE without re-training the model (training data: r = [0.55, 0.64]; test data: r = [0.58, 0.65]). Next, we performed a variance partitioning analysis to quantify the contribution of each network in predicting subjective fear experience. Synergistic signatures were developed using eight out of nine networks as well as the remaining single network, separately. Subsequently, we fitted a series of linear models to predict fear ratings using signature responses from the two models. For this analysis, we trained the model using data from randomly selected 51 participants and tested it using data from the remaining 25 participants (repeated 10,000 times). We found that each single network showed modest prediction capability (R^2^ = [0.15 0.30]), but no significant differences were found between them (FDR P > 0.5). Notably, the unique contributions of single networks were minimal (accounting for 2.89 ± 0.72% of the total R^2^; Supplementary Table 1). In summary, our results confirm the redundancy hypothesis, illustrating that subjective fear is encoded in a redundant manner, with no specific subsystem playing a crucial role in predicting fear.

## Discussion

Contemporary neuroscience models of conscious emotional experience emphasize a mental state-dependent recruitment and interaction between brain systems. However, these integrative brain models have not been empirically tested under naturalistic conditions. Here we developed a fMRI-based whole-brain model for the subjective experience of fear that capitalizes on synergistic contributions from activity and connectivity features (i.e., the CAFE). While conventional fMRI activation-based signatures failed to precisely predict fear in naturalistic contexts fusing immersive naturalistic stimuli with multivariate predictive modeling of connectivity and activity profiles allowed for the first time an accurate tracking of stable and dynamic fear experiences across naturalistic contexts, thus providing critical support for the validity of synergistic brain models under ecologically valid conditions. We further demonstrated that the CAFE – to a large extent – captures fear-specific experiences which suggests that subjective fear or the reappraisal of fear may have idiosyncratic neural representations that generalize across naturalistic contexts. Finally, we showed that subjective fear experience was represented redundantly in the whole-brain such that distributed brain regions and pathways were fear-predictive, but no system or circuit provided much unique information.

Recent studies have capitalized on multivariate pattern analysis to develop brain activation-based biomarkers for non-specific negative affect(20, 21) or fear(10, 11). These models hold the potential to advance the understanding of the brain mechanisms that establish conscious emotional experiences as well as to promote biomarker and treatment development for mental disorders characterized by exaggerated negative emotional experience(12, 14, 47, 48). However, while animal studies have demonstrated that the behavioral and physiological responses of fear are mediated by dynamic interactions between subcortical and cortical systems(5, 49–51) and that the corresponding circuit level models generalize to fear in naturalistic contexts(5, 6, 8), it remains unclear whether and how emotional experiences are represented in brain pathways and whether the neuroimaging-based brain models developed in laboratory settings are valid in real-world contexts(26).

Here, we tested for the first time the predictive capacity of these activation-based signatures on naturalistic and immersive emotional stimuli in a close-to-real-life setting(29–32). We found that the established activation-based signatures (AFSS(11), VIFS(10), PINES(20) and GNAS(21)) exhibited poor predictions across naturalistic contexts, especially for dynamic fear experience. These findings challenge the generalizability of the current and probably most precise fMRI-based brain models to conscious emotional experiences in everyday life. Against this background we developed a synergistic fMRI-based signature (CAFE) to predict fear across naturalistic contexts. The CAFE accurately predicted fear in stable as well as highly dynamic naturalistic environments which may have been facilitated by two innovations: (1) based on recent findings suggesting that neural signatures developed on functional connectivity profiles in naturalistic contexts may capture attention changes in the same naturalistic environment(35, 36) the CAFE capitalized on both, activity and connectivity features, and (2) given that the generalizability of the previous connectivity signature to novel contexts (i.e., new movies) remained limited(35), probably due to the lack of sufficient prototypical training examples or overfitting that may occur in the excessively dependent training data, the CAFE was developed on immersive video clips presenting a range of different but prototypical fear-inducting situations. Importantly, the connectivity profiles but not activation patterns critically facilitated the successful decoding of subjective fear experience in dynamic naturalistic contexts, indicating that the conscious experience of fear in everyday life may be mediated by adaptive interactions between distributed brain systems.

From the perspective of developing affective brain signatures, it is critical to control for the contribution of non-specific emotional processes such as arousal and valence which are inherently associated with the emotional experience(27, 42). In study 4 we provided evidence that the prediction performance of CAFE was generalizable across naturalistic stimuli and MRI systems and that the CAFE captured to a large extent fear-specific information. These findings suggest that the conscious experience of basic emotions such as fear is characterized by idiosyncratic neural representations or at least idiosyncratic interpretations.

This synergistic model additionally allowed us to uncover two principles underlying the architecture of subjective fear experience in naturalistic contexts. First, in contrast to prior research indicating that the behavioral or physiological response to threatening stimuli, whether conscious or unconscious, is linked to subcortical regions (for discussions see e.g., refs. 26, 52), our study found that the encoding of conscious fear experience involved the engagement of a broad set of cortical and subcortical regions as well as their interactions. This result aligns with recent appraisal(53) and constructionist(2) theories of emotion proposing that emotional experiences emerge from intricate interactions among multiple systems including core affect, sensory, memory, motor, and cognitive systems (for supporting evidence from neural decoding approaches see also refs. (10, 20, 54, 55)) as well as with recent studies on consciousness, which emphasize that conscious processing emergences from a network of interconnected brain areas distributed throughout the brain (56–58). Our findings bridge the gap between animal models that traditionally aim at translating neural circuit mechanisms of behavioral and physiological fear responses into ecologically valid contexts(5, 6, 8) and human theoretical models hypothesizing that conscious emotional experience is mediated by interactions between brain systems(1–4). Second, brain regions and pathways encoded subjective fear in a redundant fashion such that activity and connectivity characteristics of regions or large-scale networks conveyed very little unique information about subjective fear experience. Given that fear is essential for survival(59) the redundant representations may allow appropriate reactions to danger even after damage to the neural system or under suboptimal conditions. In support of this view, patients with damage to the bilateral amygdala which is traditionally considered as the ‘center’ of behavioral and physiological fear(14) can experience fear and panic in response to breathing CO2-enriched air(60).

We capitalized on recent progress in naturalistic neuroimaging and employed movies to induce and measure subjective fear under more naturalistic conditions. Movies allow a rich, multisensory and immersive experience that can model certain aspects of fear in a more naturalistic fashion than the conventional neuroimaging paradigms using isolated static stimuli. Nevertheless, this strategy cannot encompass all essential aspects of fear processing in naturalistic contexts, such as the self-relevance of the danger or active interaction with the situation. While this approach represents a substantial progression in facilitating ecologically valid neural decoders, future research should improve the approximation of real-life fear experiences, possibly through the utilization of first-person virtual reality environments or mobile neuroimaging technologies. In the current study, we identified the neural basis underlying conscious fear experience. However, the extent to which the CAFE tracks hard-wired and not necessarily conscious aspects of the fear response (e.g., behavioral activation and physiological reactivity towards subliminally presented threatening stimuli) and how conscious appraisal may itself affect the CAFE prediction will require further exploration. Moreover, it is important to note that the functional connectivity approach that used in this study can only represent un-directed associations between brain regions. How causal interactions between distributed brain regions contribute to the subjective fear experience will be an interesting avenue for future research.

In conclusion, we developed a fMRI-based synergistic signature, which capitalizes on brain activity and connectivity features and captures to a large extent fear-specific information, that accurately predicts stable and dynamic fear experience in naturalistic environments. In contrast, conventional activation-based signatures have limited ability to capture dynamic emotional experiences. Our findings further demonstrate that the neurobiology underlying subjective fear is represented by not only a broad set of brain regions but also distributed pathways in a redundant fashion. Overall, this study reveals a comprehensive functional brain architecture for subjective fear and provides an ecologically valid brain-based biomarker predictive of subjective fear intensity with high sensitivity and specificity.

## STAR Methods

### Participants

Eighty healthy, right-handed participants were recruited from Southwest University (China) for study 1. Exclusion criteria included color blindness, current or regular substance or medication use, current or history of medical or psychiatric disorders, and any contraindications for MRI. The experiments were aborted for two participants due to excessive fear, and data from another two participants were excluded from analysis due to a lack of fear experience (i.e., participants reported no fear experience during any of the videos). This resulted in a final sample of n = 76 participants (44 females; mean ± SD age = 20.21 ± 1.42).

Thirty-six participants (17 females; age = 21.03 ± 2.22) from the University of Electronic Science and Technology of China participated in study 2. Exclusion criteria included color blindness, current or regular substance or medication use, current or history of medical or psychiatric disorders, and any contraindications for MRI.

Details of the study 3 cohort were reported in previous studies(10, 11). Briefly, 31 participants (15 females; age = 23.29 ± 4.21) underwent a 1-hour fMRI session at the Institutional Review Board of Advanced Telecommunications Research Institute International (ATR), Japan.

For Study 4, a total of 80 healthy, right-handed participants from the University of Electronic Science and Technology of China were initially screened, and 63 individuals (25 females; age = 19.87 ± 2.01) were enrolled. Exclusion criteria included low arousal responses to positive or negative videos during a behavioral pre-test, color blindness, current or regular substance or medication use, current or history of medical or psychiatric disorders, and any contraindications for MRI.

In addition, a total of thirty independent participants (10 females; age = 20.87 ± 2.26) from the University of Electronic Science and Technology of China took part in the behavioral rating study for the naturalistic horror movies.

All participants provided written informed consent, and the studies were approved by the local ethics committees at Southwest University (Study 1), University of Electronic Science and Technology of China (Studies 2, 4, and the behavioral study), and the Institutional Review Board of Advanced Telecommunications Research Institute International (ATR), Japan (Study 3). The experiments were conducted in accordance with the most recent revision of the Declaration of Helsinki, and participants were compensated after the completion of the experiment.

### Naturalistic movie clips and fear rating paradigm in study 1

During the fear rating fMRI paradigm 38 short fear-inducing video clips were presented over 4 runs with each run encompassing 9 or 10 videos (including content covering humans, animals, and scenes, e.g., first view of being attacked by a snake or doing an extreme sport). Participants were instructed to attentively watch the videos and rate their level of fear experience following each video. The videos lasted 30 – 50 s followed by a 1 – 1.5 s jittered fixation-cross separating the stimuli from the rating period. During the subsequent 5 s period participants reported the level of fear they experienced for each stimulus using a 9-point Likert scale with 1 indicating no fear and 9 indicating very strong fear followed by a jittered 10 – 12 s inter-trial-interval (fixation-cross) (Fig. 1). The movie clips for the fear rating paradigm were selected from a larger database of potentially suitable video clips and selected by pre-study ratings in an independent sample (n = 30; 15 females; age = 23.23 ± 2.03). Key exclusion criteria for the movie clips were (1) a lack of fear-specificity as operationalized by more than 2 participants (∼5%) reporting an induction of non-fear negative (e.g., disgust and sad) or positive (e.g., happiness and joy) emotions during the movie, and (2) low emotional consistency over the duration of the clip, operationalized as a mean consistency rating ≤ 6.5 on a 9-point Likert scale with 1 indicating that the intensity of the elicited emotion was very inconsistent throughout the video clip and 9 indicating that the intensity of the elicited emotion was very consistent throughout the video clip.

### Naturalistic movie watching paradigm in studies 1 and 2

Following the fear rating paradigm participants in study 1 were shown a full-length short horror movie (“Don’t Look Away” by Christopher Cox, duration 7min 28 s, adapted version) with concomitant fMRI recordings. Similarly, participants in study 2 watched a segment from the horror movie “The Conjuring 2” (starting at 30 min 12 s, with a total duration of 11 min 18 s) during fMRI after they underwent a fMRI rating paradigm (data were not used in the current study). To facilitate a naturalistic and ecologically valid emotional experience no response was required from the subjects during the movie.

### Paradigm study 3

Participants were presented with 3600 images consisting of 30 animal categories and 10 object categories (90 different images per category). The stimuli were grouped in blocks of 2, 3, 4 or 6 images of the same category with each stimulus presented for 1 s (no inter-block or inter-stimulus interval). Subjective fear ratings (0 = ‘no fear’ to 5 = ‘very high fear’) for each category were established before the fMRI procedure without presenting any fearful stimuli. See also ref.(11) for the details of the paradigm.

### Dynamic arousal stimuli and arousal rating paradigm in study 4

Participants were presented with positive, negative, and neutral movie clips to induce high arousal and rated their arousal experience during each clip. A total of 40 short videos was presented over 3 runs with each run encompassing 13 or 14 videos (including content depicting humans, animals, and scenes). Stimulus presentation and subjective emotional experience reports were similar to study 1 except that participants were asked to report their arousal during each video using a 9-point Likert scale with 1 indicating very low arousal and 9 indicating very high arousal. Importantly, to avoid confounding of the stimulus materials on the decoding performance the stimuli between the studies 1 and 4 did not overlap. Videos in study 4 were selected to induce a high negative or positive arousal while neutral (low arousing) video clips were included to serve as the reference condition. The videos were initially selected based on extensive pre-study ratings (valence, arousal, and emotional category) with an independent sample (n = 26; 10 females, age = 21.62 ± 1.92) for a total of 80 videos. Valence and arousal were rated using 9-point Likert scales where 1 indicated extremely negative/very low arousal, 5 indicated neutral/medium arousal and 9 indicated extremely positive/very high arousal. Emotion category was selected from sadness, happiness, fear, anger, surprise, disgust, neutral and ambiguous. In line with Study 1, the consistency of the emotional intensity was rated on a 9-point Likert scale. The final stimulus set for the fMRI experiment included 20 low-arousal (mean arousal < 3.50), 10 high-arousal positive (mean arousal > 6.50; mean valence > 6.84) and 10 high-arousal negative (mean arousal > 6.65; mean valence < 3.31) videos. We additionally labeled 14 videos from the low-arousal stimuli as neutral stimuli (> 80% participants selected neutral for these videos), 7 videos from the high-arousal positive stimuli as happiness stimuli (> 80% participants selected happiness for these videos), 5 videos from the high-arousal negative stimuli as disgust stimuli (> 80% participants selected disgust for these videos) and 5 videos from the high-arousal negative stimuli as fear stimuli (76% participants selected fear for 1 video and > 80% selected fear for the other 4 videos). In addition. the mean ratings of the consistency of the emotional intensity were all higher than 6.5. Stimulus presentation and behavioral data acquisition were controlled using MATLAB 2014a (Mathworks, Natick, MA) and Psychtoolbox (http://psychtoolbox.org/).

### Behavioral rating paradigms

A total of thirty independent participants (10 females; age = 20.87 ± 2.26) from the University of Electronic Science and Technology of China took part in a behavioral rating study. During the experiment, participants watched the horror movie “Don’t Look Away” and the segment from the horror movie “The Conjuring 2”. Throughout the viewing, participants continuously reported their subjective fear ratings using a mouse on an incremental scale ranging from 0 to 1. The scale was labeled with “No fear” at the bottom and “Extreme fear” at the top. Additionally, markers denoting “Low fear,” “Medium fear,” and “High fear” were strategically placed at ¼, ½, and ¾ intervals along the scale, respectively. The order of the movies was counterbalanced across participants, and a 10-minute break was provided between the two movies. Stimulus presentation and behavioral data acquisition were controlled using MATLAB 2014a (Mathworks, Natick, MA) and Psychtoolbox (http://psychtoolbox.org/). Upon completion of the experiment, participants received 40 RMB for their involvement.

### MRI data acquisition and preprocessing

The details of MRI data acquisition, preprocessing, and denoising were shown in the Supplementary information.

### Predictions of previously developed activation-based signatures

We applied the AFSS, VIFS, PINES and GNAS to the naturalistic fMRI data collected in study 1 (“Don’t look away”) and study 2 (“The Conjuring 2”) to test whether these fear-related activation-based signatures could predict dynamic fear experience. Moreover, we also applied these signatures to movie clips collected in study 1 (fear experience) and study 4 (arousal experience) to test the sensitivity and specificity of the signatures on dynamic stimuli separately. In line with preprocessing pipelines used in refs. (10, 20) in these analyses we used SPM12 preprocessed data.

### Fear signature development

The sample in study 1 was split into training (n = 51; i.e., 2/3 of the total participants) and test (n = 25) datasets. Next, we developed multivariate fear signatures through whole-brain activation-based (incorporating activation for each voxel; n = 228,022 features), functional connectivity-based (including connectivity between each pair of parcels; n = 106,953 features), and synergistic (integrating mean activation for each parcel and functional connectivity between each pair of parcels; n = 107,416 features) approaches using the training data only (n = 341 samples). Following our previous studies(10, 25), we applied the linear SVR algorithm with the cost parameter C = 1 and epsilon = 0.01 to develop the fear signatures. This algorithm was implemented in the CanlabCore toolbox (https://github.com/canlab/CanlabCore), which utilizes the Spider toolbox (http://people.kyb.tuebingen.mpg.de/spider). The parcellation used to develop connectivity-based and synergistic signatures included 400 cortical regions from the Schaefer atlas(40), 34 subcortical regions from the Melbourne subcortex atlas (n = 32)(61) and the reinforcement learning atlas (extended amygdala and hypothalamus)(62), and periaqueductal gray (PAG), brainstem and cerebellar regions from ref.(34) (n = 29), resulting in a total of 463 regions. Additionally, another parcellation consisting of 360 cortical regions, 63 subcortical and cerebellar regions, as well as PAG and brainstem regions(34), was used to validate our findings.

To evaluate model performance, we assessed the prediction-outcome correlation and coefficient of determination (R^2^) within the training data using cross-validated analysis (10 repeats of 10-fold cross-validation), as recommended by refs.(63, 64). R^2^ represents the percentage of fear rating variance accounted for by the prediction models, and it is calculated using the following formula::

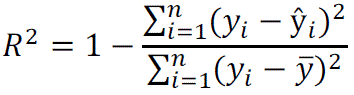

where y_i_ is the true fear rating for the i-th sample, 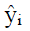 is the model predicted rating for the i-th sample, and 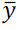 is the mean of all fear ratings. To avoid the potential bias in train-test split, we re-ran this procedure 100 times.

In addition to cross-validated performance in the training data, we evaluated the performance of the synergistic signature (i.e., the CAFE) for predicting fear in the independent test data. Significance testing was conducted using one-tailed permutation test: P = (1+number of r/R^2^ >= empirical r/R^2^)/(1+number of permutations).

### Prediction of subjective fear during naturalistic contexts (movie)

To test the ecological validity of the fear-related signatures under naturalistic conditions in terms of predicting subjective fear during naturalistic movie watching (without any response) we calculated signature response using a tapered sliding window approach, where the parcel mean activation and connectivity among parcels were computed and the dot product of the vectorized feature with signature weights was employed within the temporal window. We implemented a window size of 22 TR (= 44 s) for both “Don’t Look Away” and “The Conjuring 2” data, following the optimal window size suggested by previous literature, with a step size of 1 TR and a Gaussian kernel σ = 3 TR(35, 38). To better evaluate the prediction capabilities of the activation-based signatures, in addition to utilizing the sliding window approach, we also directly computed the signature response for each individual TR. Since we were interested in whether the model could predict temporal dynamics, unlike previous studies(35, 36), we did not convolve fear rating with the HRF or slid fear rating. Given that we were interested in whether the model captures temporal dynamics rather than the actual values of group-average fear ratings we employed correlation as a metric to assess the predictive performance of our model. P values were obtained from permutation tests, where observed correlation coefficient and R^2^ were compared with null distributions of correlation coefficient between signature response and 10,000 permuted, phase-randomized fear ratings (one-tailed).

### Sensitivity and specificity of the CAFE

We applied the CAFE to fMRI data collected in study 4 using a dot product of each vectorized feature with signature weights to determine how much the signature depend on arousal negative valence features separately. The signature response served as the basis for single-interval classification (e.g., fear vs. disgust) with a cutoff selected to maximize overall accuracy. To better match arousal ratings between the emotional categories, we excluded data from non-neutral movie clips that induced only low-to-medium levels of subjective arousal (i.e., arousal ≤ 6 on a 9-point Likert scale). This allowed to generate a dataset with comparably high arousal induction for fear and disgust (fear: 7.91 ± 0.57; disgust: 7.78 ± 0.54; P = 0.17, BF_01_ = 2.83), although the fear clips still induced slightly higher arousal than the happiness clips (7.56 ± 0.49) (P = 1.00 × 10^-4^). Bayesian factor analyses (https://richarddmorey.github.io/BayesFactor/) were used to compare the likelihood of the data fitting under the null hypothesis with the likelihood of fitting under the alternative hypothesis. BF_01_ represents the ratio that contrasts the likelihood of the data fitting under the null hypothesis with the likelihood of fitting under the alternative hypothesis. In contrast, BF_10_ represents the ratio that contrasts the likelihood of the data fitting under the alternative hypothesis with the likelihood of fitting under the null hypothesis.

### Determining subjective fear representation

To test which brain regions and circuits reliably encode subjective fear information we constructed 10,000 randomly selected samples from study 1 consisting of paired brain (activation and connectivity features) and outcome data and ran SVR on each. Two-tailed, uncorrected P-values were calculated for each feature based on the proportion of SVR weights below or above zero and subjected to false discovery rate (FDR) correction. To further explore the neurobiology of subjective fear we ran parcel-based prediction analysis and calculated the model encoding maps using the following formula(44): A = cov(X) × W × cov(S)^-1^, where A is the reconstructed modeling encoding map, cov(X) is the covariance matrix of training data, W is the weights of the SVR model, and cov(S) is the covariance matrix of the latent factors (i.e., model predictions), which is defined as W^T^ × X. All statistical maps were thresholded at FDR P < 0.05.

To further test how fear intensity information is encoded in brain systems, we conducted a series of experiments. To this end we divided the brain into 9 networks, including 7 resting-state (65), 1 subcortical and 1 cerebellar (including PAG and brainstem)(34) network. We then trained models with a range of randomly selected regions (40–400) from various networks and evaluated their performances (repeated 1,000 times). Next, we conducted “variance partitioning” and “virtual lesion” analyses. Specifically, synergistic signatures were developed using 8 out of 9 networks, which included the activation and connectivity among parcels in those networks, as well as the remaining single network, which included the mean activation of each parcel in the network and all functional connectivity with parcels within that network, separately. Subsequently, we fitted a series of linear models to predict fear using signature response from the two models. This analysis was repeated 9 times until each network being the remaining 1 network once. We then performed a variance partitioning analysis to decompose the proportion of explained variance to the variance explained only by the 8-network-model predictions (ΔR^2^_8net_), variance explained only by the 1-network-model predictions (ΔR^2^_1net_), and variance explained by predictions from both model trained on both 8 networks and model trained on the remaining 1 network (R^2^_8net∩1net_). For this analysis, we trained the model using data from 51 participants and tested it using data from the remaining 25 participants. To mitigate potential bias in the train-test split, we repeated this procedure 10,000 times.

## Acknowledgements

The study was supported by the National Natural Science Foundation (NSFC, 32250610208; 82271583; 32300862), the National Key Research and Development Program (2018YFA0701400) of China, the Ministry of Science and Technology (Brain Project 2022ZD0208500, MOST2030), the Fundamental Research Funds for the Central Universities (SWU2309733) and the Innovation Research 2035 Pilot Plan of Southwest University (SWUPilotPlan006).

## Author contributions

F.Z. and B.B. conceived and designed the experiment, interpreted the results, and wrote the manuscript. F.Z. and R.Z. collected the data. R.Z. preprocessed the data. F.Z. analyzed the data and created the figures. All authors revised the manuscript.

## Data availability

Brain patterns and source data for figures will be shared upon publication through a GitHub repository. Raw fMRI data will be shared in the OpenfMRI upon publication.

## Code availability

The code for generating the figures and main analyses will be shared upon publication through a GitHub repository.

## Supplementary information for

**This file includes:**

**Supplementary methods**

**Supplementary results**

**Supplementary Figures 1 to 5**

**Supplementary Table 1**

**Supplementary references**

## Supplementary Methods

### MRI data acquisition in Study 1

MRI data were collected on a 3.0-T SIEMENS PRISMA Fit MRI system (Erlangen, Germany). Functional MRI data was acquired using a T2*-weighted echo-planar imaging (EPI) pulse sequence (repetition time = 2000 ms, echo time = 30 ms, flip angle = 90°, number of slices = 62, slice orientation = transversal, posterior to anterior phase encoding, voxel size = 2 mm isotropic, gap between slices = 0 mm, field of view = 224 × 224 mm, iPAT (GRAPPA) factor = 2, echo spacing = 0.54 ms). To improve spatial normalization and exclude participants with apparent brain pathologies high-resolution T1-weighted images were acquired using a three-dimension (3D) magnetization-prepared rapid gradient-echo (MP-RAGE) sequence (repetition time = 2530 ms, echo time = 2.98 ms, 1 × 0.5 × 0.5 mm voxels).

### MRI data acquisition in study 2

MRI data were collected on a 3.0-T GE Discovery MR750 system (General Electric Medical System, Milwaukee, WI, USA). Functional MRI data was acquired using a T2*-weighted echo-planar imaging (EPI) pulse sequence (repetition time = 2000 ms, echo time = 30 ms, 36 slices, slice thickness = 3.8 mm, no gap, field of view = 200 × 200 mm, resolution = 64 × 64, flip angle = 90°, 3.125 × 3.125 × 3.8 mm voxels, posterior to anterior phase encoding). To improve spatial normalization and exclude participants with apparent brain pathologies a high-resolution T1-weighted image was acquired using a 3D spoiled gradient recalled (SPGR) sequence (176 slices, repetition time = 8.22 ms, echo time = 3.15 ms, field of view = 256 × 256 mm, resolution = 256 × 256, flip angle = 8°, 1 × 1 × 1 mm voxels).

### MRI data acquisition in study 3

Participants were scanned in two 3T MRI scanners (Prisma Siemens and Verio Siemens) at the ATR Brain Activation Imaging Center. Functional MRI data was acquired using an EPI pulse sequence (repetition time = 2000 ms, echo time = 30 ms, 33 slices, slice thickness = 3.5 mm, no gap, field of view = 192 × 192 mm, resolution = 64 × 64, flip angle = 80°, voxel size = 3 × 3 × 3.5 mm). T1-weighted image was acquired using a three-dimension (3D) magnetization-prepared rapid gradient-echo (MP-RAGE) sequence (256 slices, TR = 2250 ms, TE = 3.06 ms, voxel size = 1 × 1 × 1 mm, field-of-view = 256 × 256 mm, matrix size = 256 × 256, slice thickness = 1 mm, 0 mm slice gap, TI = 900 ms, flip angle = 9°. The least-square separate single-trial analysis approach was employed to iteratively fit a GLM to estimate the brain response to the first image of each block and then the within-subject beta images with the same fear ratings were averaged, which resulted in one beta map per rating for each subject (for paradigm, MRI acquisition and analysis details see ref.(1)).

### MRI data acquisition in study 4

MRI data were collected on a 3.0-T GE Discovery MR750 system (General Electric Medical System, Milwaukee, WI, USA). Functional MRI data was acquired using a T2*-weighted echo-planar imaging (EPI) pulse sequence (repetition time = 2000 ms, echo time = 30 ms, 39 slices, slice thickness = 3mm, gap between slice = 1 mm, field of view = 240 × 240 mm, resolution = 64 × 64, flip angle = 90°, 3.75 × 3.75 × 4 mm voxels, posterior to anterior phase encoding). To improve spatial normalization and exclude participants with apparent brain pathologies a high-resolution T1-weighted image was acquired using a 3D spoiled gradient recalled (SPGR) sequence (repetition time = 5.96 ms, echo time = 1.97 ms, 1 × 1 × 1 mm voxels).

### fMRI data preprocessing

Structural and functional MRI data in studies 1, 2 and 4 were preprocessed using FMRIPREP 21.0.0(2) (RRID: SCR_016216), a Nipype 1.6.1(3) based tool that integrates preprocessing routines from different software packages. Spatial normalization to the ICBM 152 Nonlinear Asymmetrical template version 2009c was performed through nonlinear registration with the antsRegistration tool of ANTs v2.3.3(4), using brain-extracted versions of both T1w volume and template. Brain tissue segmentation of cerebrospinal fluid (CSF), white matter (WM) and gray matter (GM) was performed on the brain-extracted T1w using FAST (FSL v6.0.5.1)(5). Before the automated preprocessing, 4 initial volumes of fMRI data were removed in order to allow for image intensity stabilization. Next, functional data was slice time corrected using 3dTshift from AFNI(6) and motion corrected using mcflirt (FSL v6.0.5.1). This was followed by co-registration to the corresponding T1w using boundary-based registration(7) with six degrees of freedom, using FLIRT (FSL). Motion correcting transformations, BOLD-to-T1w transformation and T1w-to-template (MNI) warp (interpolated to 2 mm isotropic voxels) were concatenated and applied in a single step using ants ApplyTransforms (ANTs v2.3.3) using Lanczos interpolation.

To facilitate a fair comparison of the prediction performance between the developed synergistic signature and previous developed activation-based signatures(1, 8–10) we additionally preprocessed fMRI data using SPM 12 (v7771; https://www.fil.ion.ucl.ac.uk/spm/software/spm12/) following the preprocessing protocols in our previous study(8). Briefly, functional images were corrected for differences in the acquisition timing of each slice and spatially realigned to the first volume and unwarped to correct for nonlinear distortions related to head motion or magnetic field inhomogeneity. The anatomical image was segmented into grey matter, white matter, cerebrospinal fluid, bone, fat and air by registering tissue types to tissue probability maps. Next, the skull-stripped and bias corrected structural image was generated and the functional images were co-registered to this image. The functional images were subsequently normalized the MNI space (interpolated to 2 × 2 × 2mm voxel size) by applying the forward deformation parameters that were obtained from the segmentation procedure, and spatially smoothed using an 8-mm full-width at half maximum (FWHM) Gaussian kernel.

To perform denoising, we fit a voxel-wise general linear model (GLM) for each participant. Specifically, in line with ref. (11), we removed variance associated with the mean, linear and quadratic trends, the average signal within anatomically-derived CSF mask, the effects of motion estimated during the head-motion correction using an expanded set of 24 motion parameters (six realignment parameters, their squares, their derivatives, and their squared derivatives) and motion spikes (FMRIPREP default: FD > 0.5mm or standardized DVARS > 1.5)(12). For the fear and arousal rating data we additionally constructed a task-related regressor with the rating period convolved with the canonical hemodynamic response function (HRF).

For the details of preprocessing and GLM analysis for study 3 please see ref. (1).

### Data averaging

To develop the signatures predictive of subjective fear we first generated subjective fear-related activation and functional connectivity features for each subject. To this end, for each participant, we concatenated the fMRI data with the same fear level rating during the video presentation period. Of note, the onset time for each video was shifted by 2-3 TRs because of the hemodynamic lag. To obtain relatively robust functional connectivity between regions for each fear level the concatenated data for a fear level with less than 25 time points (i.e., less than 2 videos) was excluded. We next calculated voxel-wise and parcel-wise mean activation as well as functional connectivity (Fisher z-transformed Pearson correlation coefficient) among parcels for each fear level and each subject separately. For the fMRI data collected in study 2 we employed the same method to calculate parcel-wise mean activation as well as functional connectivity (Fisher z-transformed) among parcels for each emotion and each subject separately.

## Supplementary Results

### Previously developed activation-based affective signatures cannot capture fear changes during naturalistic movie watching

Correlating the group mean signature responses of each TR with the group average fear rating (convolved with hemodynamic response function) revealed that neither AFSS (study 1: r = 0.02, permutation P = 0.374; study 2: r = –0.05, permutation P = 0.845), VIFS (study 1: r = 0.07, permutation P = 0.097; study 2: r = 0.05, permutation P = 0.125), PINES (study 1: r = 0.01, permutation P = 0.523; study 2: –r = 0.04, permutation P = 0.858), nor GNAS (study 1: r = –0.09, permutation P = 0.983; study 2: r = 0.01, permutation P = 0.479) were able to predict changes in subjective fear under naturalistic conditions. Similar findings were observed when correlating the mean signature response with the mean unconvolved fear rating (study 1: maximum r = 0.07, permutation P = 0.092 (VIFS); study 2: maximum r = 0.06, permutation P = 0.056 (VIFS)).

We further investigated whether applying these activation-based affective signatures to the naturalistic data using the tapered sliding window approach (window size = 44 s) could predict dynamic fear ratings. However, neither AFSS (study 1: r = 0.06, permutation P = 0.428; study 2: r = –0.08, permutation P = 0.680), VIFS (study 1: r = 0.13, permutation P = 0.242; study 2: r = 0.18, permutation P = 0.114), PINES (study 1: r = –0.07, permutation P = 0.556; study 2: r = –0.14, permutation P = 0.904), nor GNAS (study 1: r = –0.23, permutation P = 0.927; study 2: r = –0.02, permutation P = 0.534) were able to predict convolved subjective fear ratings. Similarly, the mean signature response of the group did not correlate with the mean unconvolved fear rating (study 1: maximum r = 0.09, permutation P = 0.312 (VIFS); study 2: maximum r = 0.18, permutation P = 0.115 (VIFS)).

### The novel synergistic signatures can better predict short-term stable fear experience

To assess whether synergistic signatures (i.e., the CAFE) improves predictions of short-term homogeneous fear experiences compared to conventional activation and connectivity signatures, we randomly split participants in study 1 into a training sample (n = 51) and a test sample (n = 25) for 100 iterations. Our findings reveal that when utilizing a parcellation of n = 425 brain regions, the performance of the synergistic signature slightly decreased (training data: mean ± standard deviation R^2^ = 0.36 ± 0.03; test data: R^2^ = 0.37 ± 0.06) compared to using a parcellation of n = 463 regions (training data: R^2^ = 0.40 ± 0.03; test data: R^2^ = 0.41 ± 0.05). Nonetheless, both versions of the synergistic signature demonstrated superior prediction performance in comparison to activation-based (training data: R^2^ = 0.33 ± 0.03; test data: R^2^ = 0.33 ± 0.05) and connectivity-based (training data: R^2^ = 0.31 ± 0.04; test data: R^2^ = 0.33 ± 0.07) signatures. These findings suggest that the CAFE method is more effective in capturing the stable subjective experience of fear.

### The VIFS and GNAS, but not the AFSS or PINES, predict video clip-induced short-term stable fear experience

We found that the VIFS (r = 0.26, P = 2×10^-9^) and GNAS (r = 0.15, P = 5×10^-4^), but not AFSS (r = –0.06, P = 0.215) or PINES (r = 0.02, P = 0.706), significantly predicted movie clip-induced fear (Supplementary Fig. 1).

Given that the CAFE was developed on the training data in study 1, we additionally applied these signatures to the hold-out test data in study 1 to facilitate a fair comparison. We found similar prediction performances (VIFS: r = 0.35, P = 5×10^-11^; GNAS: r = 0.18, P = 0.001; AFSS: r = –0.02, P = 0.676; PINES: r = 0.02, P = 0.774) as compared to the whole sample. Importantly, these prediction performances were substantially lower than that of the CAFE (r = 0.67; all permutation P of the differences were less than 1 × 10^-4^).

**Supplementary Fig. 1.**
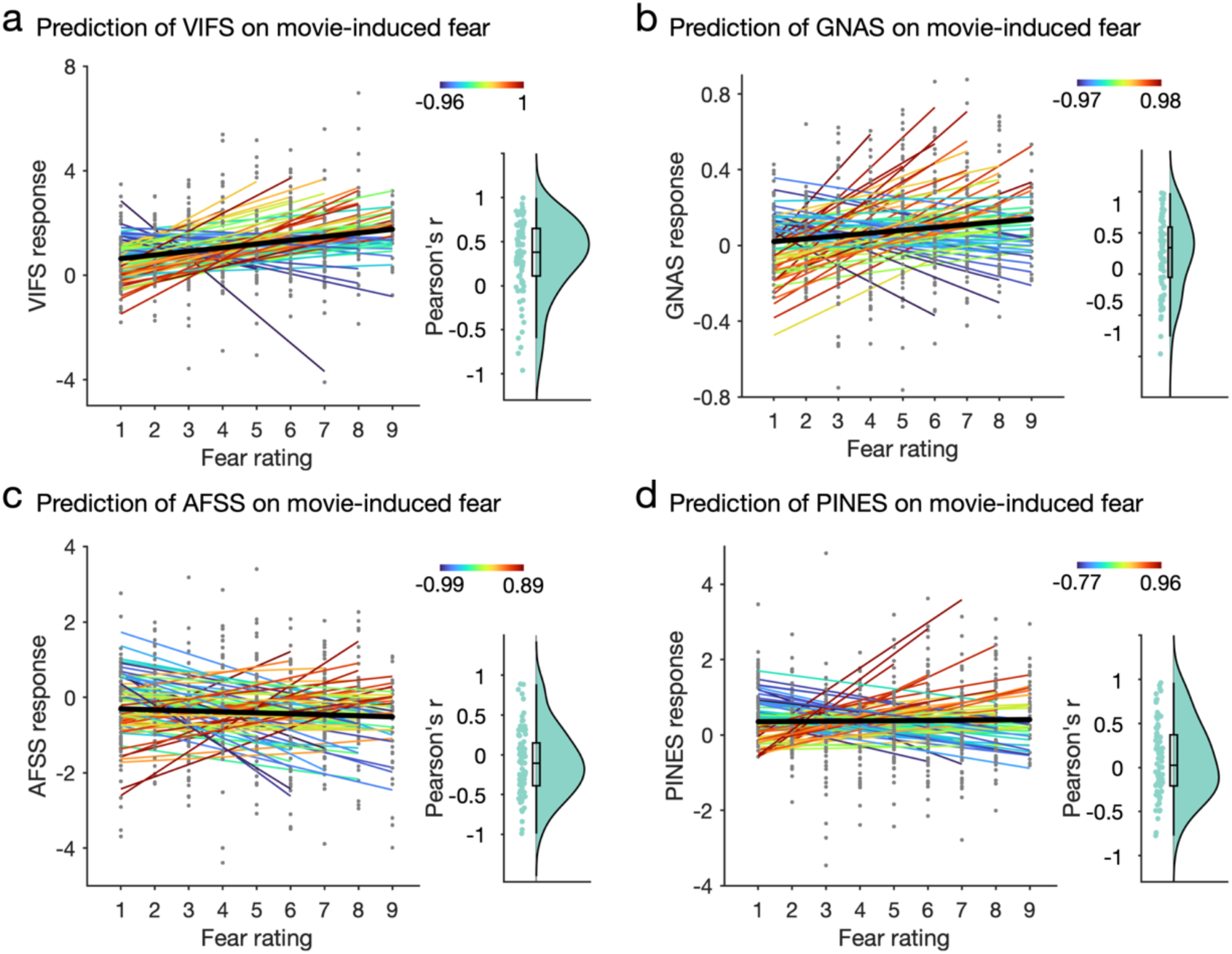
Predicting movie clip-watching induced subjective fear experiences (study 1) with previously established activation-based signatures. The VIFS (panel a; r = 0.26, P = 2×10^-9^) and GNAS (panel b; r = 0.15, P = 5×10^-4^) significantly predict movie clip-induced subjective feelings of fear whereas the AFSS (panel c; r = –0.06, P = 0.215)) and PINES (panel d; r = 0.02, P = 0.706) fail to predict movie-included subjective fear experience. VIFS, visually induced fear signature; GNAS, Generalized negative affect signature; AFSS, animal fear schema signature; PINES, picture-induced negative emotion signature.

**Supplementary Fig. 2.**
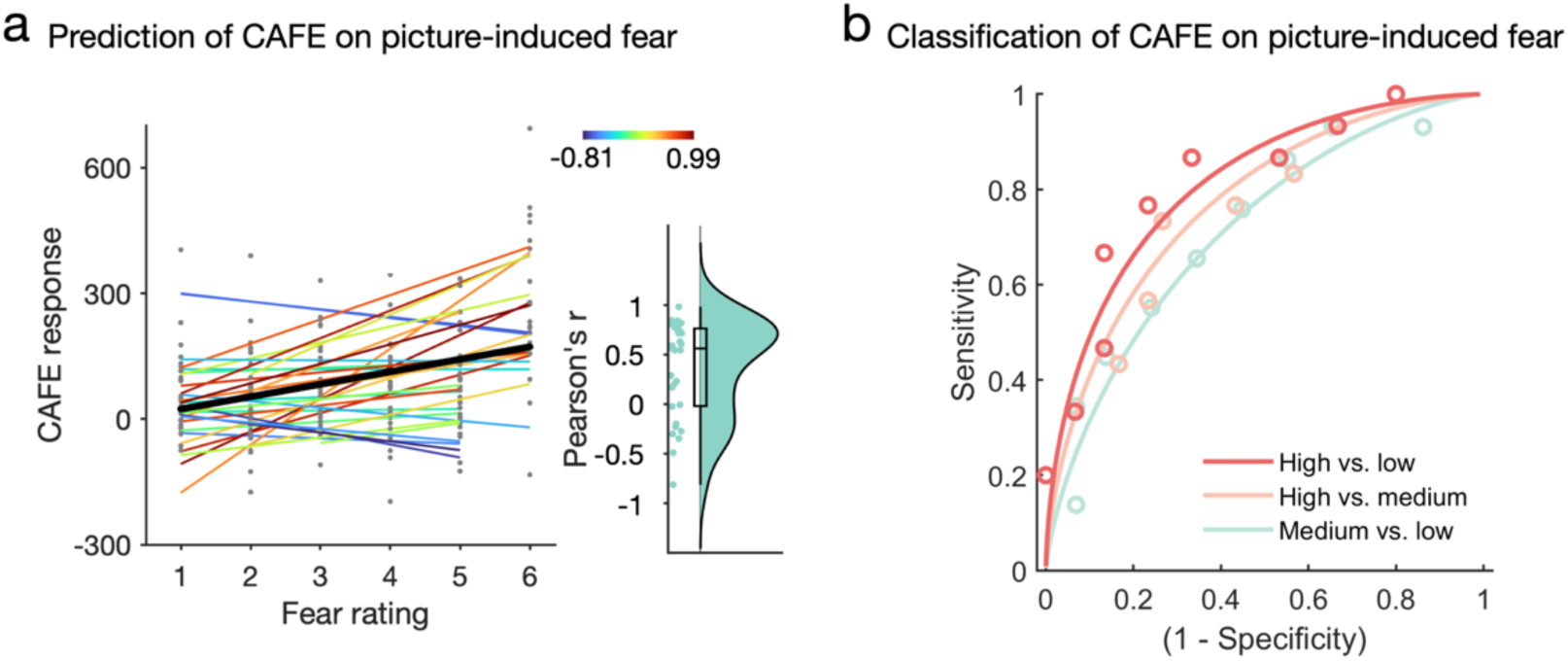
Predicting picture-induced subjective fear experiences with the activation pattern of the CAFE. a, the activation pattern of the CAFE accurately predicts subjective fear experience induced by animal pictures (overall prediction-outcome r = 0.35, P = 6×10^-6^). Each colored line represents prediction within each individual participant. The black line indicates the overall (i.e., within and between participants) prediction. Raincloud plots show the distribution of within-participant predictions. b, the activation pattern of the CAFE can classify high fear (average of rating 4 and 5) from medium fear (average of rating 2 and 3; accuracy = 73%, P = 0.016, d = 0.74) and low fear (average of rating 0 and 1; accuracy = 77%, P = 0.005, d = 0.89). In addition, the CAFE to some extent distinguishes medium fear vs. low fear (d = 0.56) although the accuracy is not significantly higher than chance level (accuracy = 66%, P = 0.136).

**Supplementary Fig. 3.**
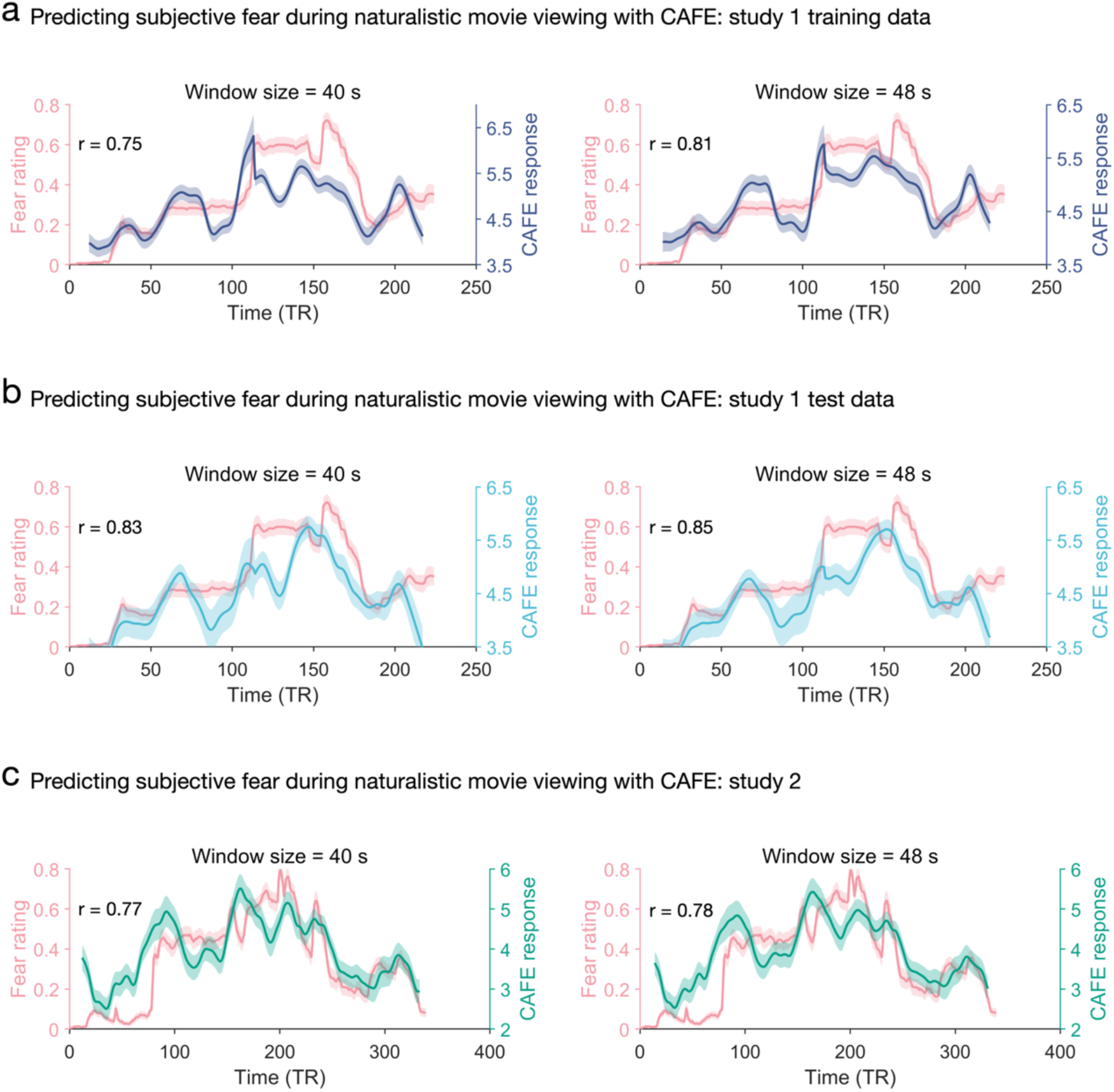
Results of group-average subjective fear prediction from the CAFE using different window sizes. The CAFE accurately tracks the dynamic fear changes using window sizes of 20 and 24 TRs in study 1 training dataset (a) and test dataset (b) as well as study 2 (c). These results suggest that our findings remain robust across selection of window sizes.

**Supplementary Fig. 4.**
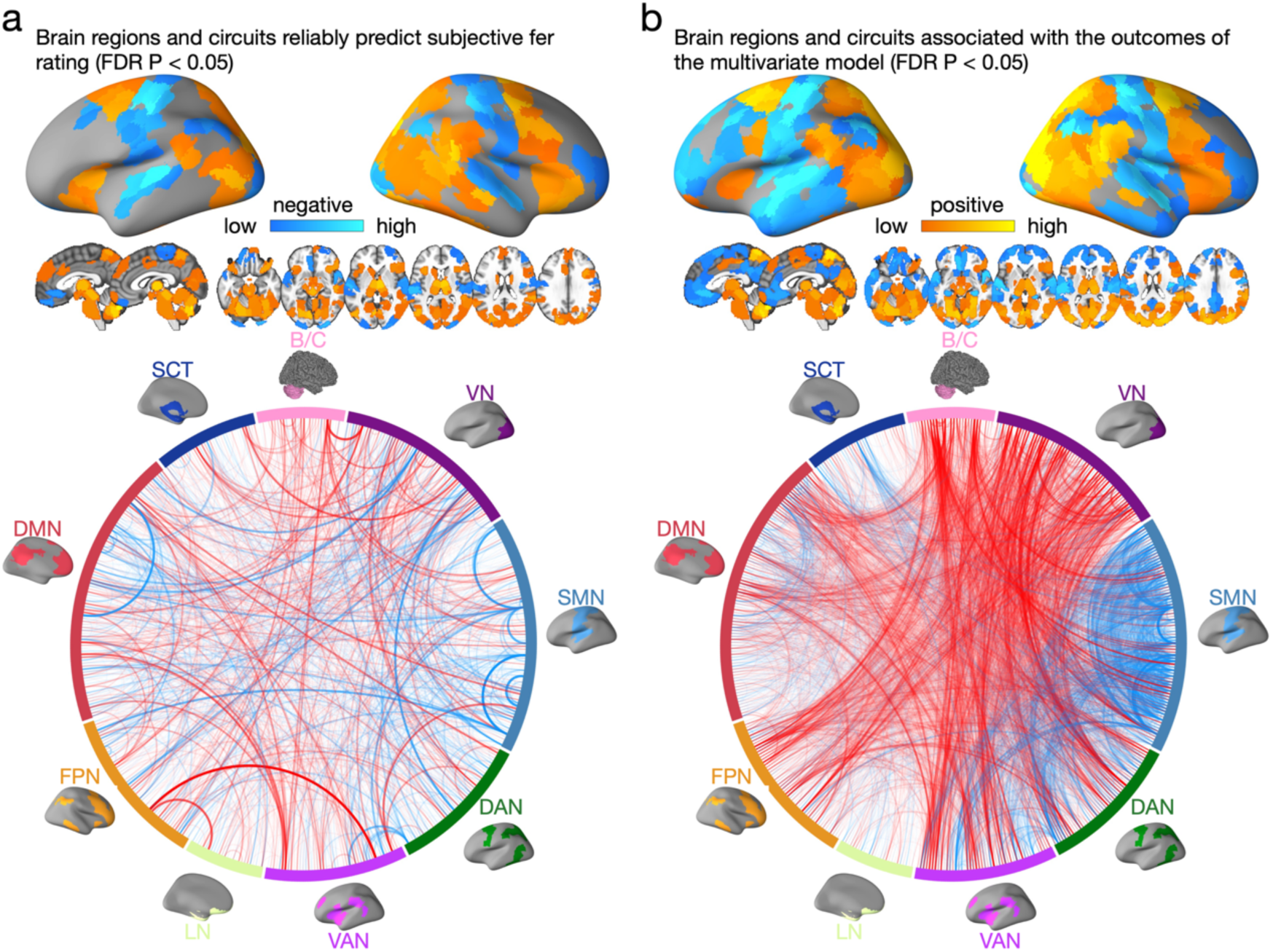
Brain regions and pathways associated with and predictive of subjective fear rating. a, model weight maps showing brain regions and circuits reliably predictive of fear ratings (FDR P < 0.05). Circular plots represent the significant pathways. b, model encoding maps showing brain regions and circuits reliably associated with the outcomes of the multivariate model (FDR P < 0.05). Circular plots represent the significant pathways.

**Supplementary Fig. 5.**
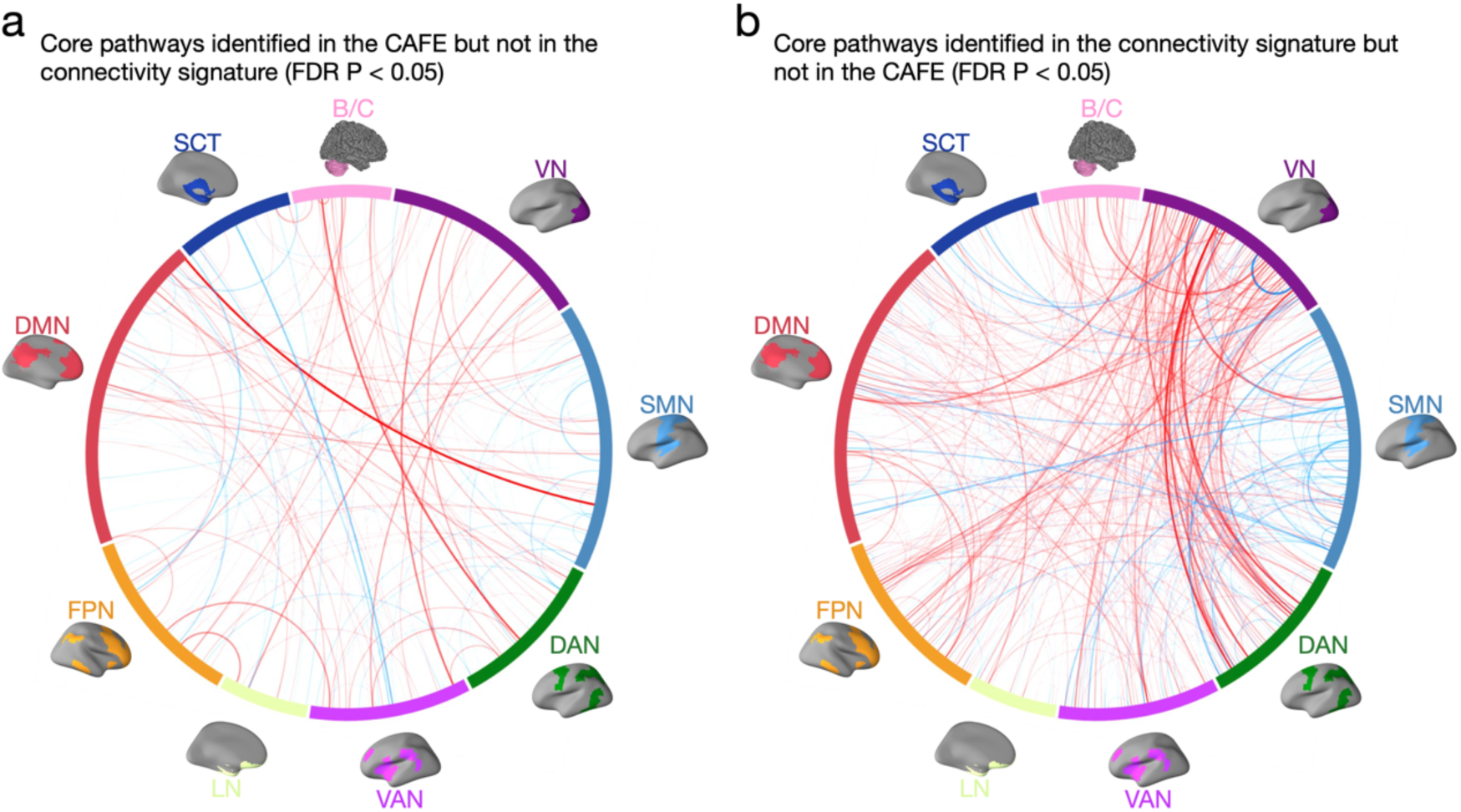
Core brain pathways for the CAFE and the connectivity signature. Circular plot visualizations of core pathways identified in the CAFE but not in the connectivity signature (panel a) and pathways identified in the connectivity signature but not in the CAFE (panel b); red line indicates positive weight and association and blue line indicates negative weight and association (all FDR P < 0.05).

**Supplementary Table 1.**
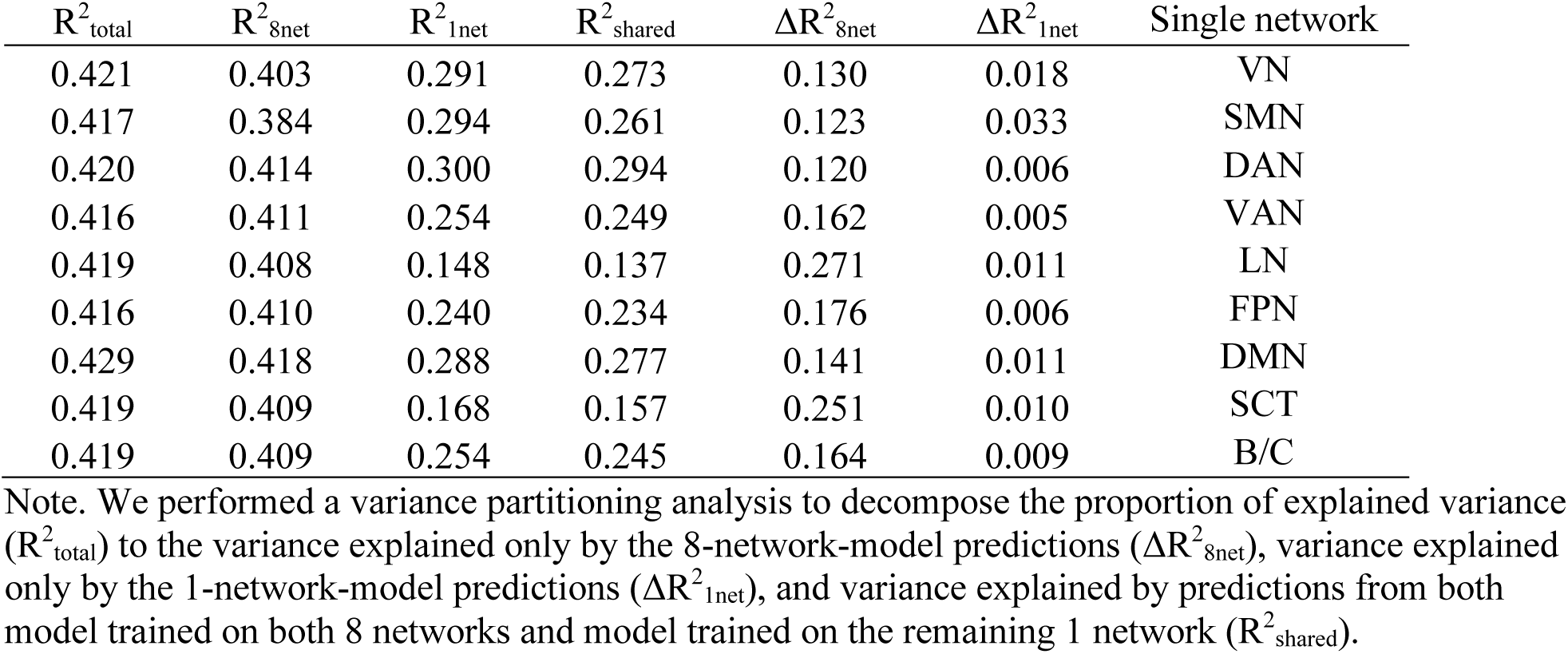
Predictive capacity of each large-scale brain network.

## References

1. R. Adolphs, How should neuroscience study emotions? by distinguishing emotion states, concepts, and experiences. Social Cognitive and Affective Neuroscience 12, 24–31 (2016).

2. L. F. Barrett, The theory of constructed emotion: an active inference account of interoception and categorization. Social Cognitive and Affective Neuroscience 12, 1–23 (2016).

3. L. Pessoa, A Network Model of the Emotional Brain. Trends in Cognitive Sciences 21, 357–371 (2017).

4. J. E. LeDoux, Thoughtful feelings. Current Biology 30, R619–R623 (2020).

5. P. H. Janak, K. M. Tye, From circuits to behaviour in the amygdala. Nature 517, 284–292 (2015).

6. P. Tovote, J. P. Fadok, A. Lüthi, Neuronal circuits for fear and anxiety. Nature Reviews Neuroscience 16, 317–331 (2015).

7. D. Mobbs, D. B. Headley, W. Ding, P. Dayan, Space, Time, and Fear: Survival Computations along Defensive Circuits. Trends in Cognitive Sciences 24, 228–241 (2020).

8. M. Zelikowsky, S. Hersman, M. K. Chawla, C. A. Barnes, M. S. Fanselow, Neuronal Ensembles in Amygdala, Hippocampus, and Prefrontal Cortex Track Differential Components of Contextual Fear. The Journal of Neuroscience 34, 8462–8466 (2014).

9. B. A. Pellman, J. J. Kim, What Can Ethobehavioral Studies Tell Us about the Brain’s Fear System? Trends in Neurosciences 39, 420–431 (2016).

10. F. Zhou et al., A distributed fMRI-based signature for the subjective experience of fear. Nature Communications 12, 6643 (2021).

11. V. Taschereau-Dumouchel, M. Kawato, H. Lau, Multivoxel pattern analysis reveals dissociations between subjective fear and its physiological correlates. Molecular Psychiatry 25, 2342–2354 (2020).

12. V. Taschereau-Dumouchel, M. Michel, H. Lau, S. G. Hofmann, J. E. LeDoux, Putting the “mental” back in “mental disorders”: a perspective from research on fear and anxiety. Molecular Psychiatry 27, 1322–1330 (2022).

13. J. E. LeDoux, S. G. Hofmann, The subjective experience of emotion: a fearful view. Current Opinion in Behavioral Sciences 19, 67–72 (2018).

14. J. E. LeDoux, D. S. Pine, Using Neuroscience to Help Understand Fear and Anxiety: A Two-System Framework. American Journal of Psychiatry 173, 1083–1093 (2016).

15. J. D. Cohen et al., Computational approaches to fMRI analysis. Nature Neuroscience 20, 304–313 (2017).

16. P. A. Kragel et al., A human colliculus-pulvinar-amygdala pathway encodes negative emotion. Neuron 109, 2404–2412.e2405 (2021).

17. R. A. Poldrack et al., Scanning the horizon: towards transparent and reproducible neuroimaging research. Nature Reviews Neuroscience 18, 115–126 (2017).

18. L. Kohoutová et al., Individual variability in brain representations of pain. Nature Neuroscience 25, 749–759 (2022).

19. T. D. Wager et al., An fMRI-Based Neurologic Signature of Physical Pain. New England Journal of Medicine 368, 1388–1397 (2013).

20. L. J. Chang, P. J. Gianaros, S. B. Manuck, A. Krishnan, T. D. Wager, A Sensitive and Specific Neural Signature for Picture-Induced Negative Affect. PLOS Biology 13, e1002180 (2015).

21. M. Čeko, P. A. Kragel, C.-W. Woo, M. López-Solà, T. D. Wager, Common and stimulus-type-specific brain representations of negative affect. Nature Neuroscience 25, 760–770 (2022).

22. P. A. Kragel, L. Koban, L. F. Barrett, T. D. Wager, Representation, Pattern Information, and Brain Signatures: From Neurons to Neuroimaging. Neuron 99, 257–273 (2018).

23. C.-W. Woo, L. J. Chang, M. A. Lindquist, T. D. Wager, Building better biomarkers: brain models in translational neuroimaging. Nature Neuroscience 20, 365–377 (2017).

24. F. Zhou et al., Empathic pain evoked by sensory and emotional-communicative cues share common and process-specific neural representations. eLife 9, e56929 (2020).

25. X. Gan et al., Rotten to the core – a neurofunctional signature of subjective core disgust generalizes to socio-moral contexts. bioRxiv 10.1101/2023.05.18.541259, 2023.2005.2018.541259 (2023).

26. D. Mobbs et al., Viewpoints: Approaches to defining and investigating fear. Nature Neuroscience 22, 1205–1216 (2019).

27. P. A. Kragel, K. S. LaBar, Decoding the Nature of Emotion in the Brain. Trends in Cognitive Sciences 20, 444–455 (2016).

28. E. Bullmore, O. Sporns, Complex brain networks: graph theoretical analysis of structural and functional systems. Nature Reviews Neuroscience 10, 186–198 (2009).

29. S. Sonkusare, M. Breakspear, C. Guo, Naturalistic Stimuli in Neuroscience: Critically Acclaimed. Trends in Cognitive Sciences 23, 699–714 (2019).

30. I. P. Jääskeläinen, M. Sams, E. Glerean, J. Ahveninen, Movies and narratives as naturalistic stimuli in neuroimaging. NeuroImage 224, 117445 (2021).

31. S. B. Eickhoff, M. Milham, T. Vanderwal, Towards clinical applications of movie fMRI. NeuroImage 217, 116860 (2020).

32. M. Khosla, G. H. Ngo, K. Jamison, A. Kuceyeski, M. R. Sabuncu, Cortical response to naturalistic stimuli is largely predictable with deep neural networks. Science Advances 7, eabe7547 (2021).

33. H. Saarimäki et al., Classification of emotion categories based on functional connectivity patterns of the human brain. NeuroImage 247, 118800 (2022).

34. J.-J. Lee et al., A neuroimaging biomarker for sustained experimental and clinical pain. Nature Medicine 27, 174–182 (2021).

35. H. Song, E. S. Finn, M. D. Rosenberg, Neural signatures of attentional engagement during narratives and its consequences for event memory. Proceedings of the National Academy of Sciences 118, e2021905118 (2021).

36. H. Song, B.-y. Park, H. Park, W. M. Shim, Cognitive and Neural State Dynamics of Narrative Comprehension. The Journal of Neuroscience 41, 8972–8990 (2021).

37. A. B. Satpute et al., Involvement of Sensory Regions in Affective Experience: A Meta-Analysis. Frontiers in Psychology 6 (2015).

38. E. A. Allen et al., Tracking Whole-Brain Connectivity Dynamics in the Resting State. Cerebral Cortex 24, 663–676 (2014).

39. D. C. Van Essen, M. F. Glasser, D. L. Dierker, J. Harwell, T. Coalson, Parcellations and Hemispheric Asymmetries of Human Cerebral Cortex Analyzed on Surface-Based Atlases. Cerebral Cortex 22, 2241–2262 (2011).

40. A. Schaefer et al., Local-Global Parcellation of the Human Cerebral Cortex from Intrinsic Functional Connectivity MRI. Cerebral Cortex 28, 3095–3114 (2017).

41. L. F. Barrett, Are Emotions Natural Kinds? Perspectives on Psychological Science 1, 28–58 (2006).

42. J. A. Russell, A circumplex model of affect. Journal of Personality and Social Psychology 39, 1161–1178 (1980).

43. L. Kohoutová et al., Toward a unified framework for interpreting machine-learning models in neuroimaging. Nature Protocols 15, 1399–1435 (2020).

44. S. Haufe et al., On the interpretation of weight vectors of linear models in multivariate neuroimaging. NeuroImage 87, 96–110 (2014).

45. A. R. Damasio et al., Subcortical and cortical brain activity during the feeling of self-generated emotions. Nature Neuroscience 3, 1049–1056 (2000).

46. N. Kriegeskorte, P. K. Douglas, Interpreting encoding and decoding models. Current Opinion in Neurobiology 55, 167–179 (2019).

47. F. Zhou et al., Human Extinction Learning Is Accelerated by an Angiotensin Antagonist via Ventromedial Prefrontal Cortex and Its Connections With Basolateral Amygdala. Biological Psychiatry 86, 910–920 (2019).

48. R. Zhang et al., Angiotensin II Regulates the Neural Expression of Subjective Fear in Humans: A Precision Pharmaco-Neuroimaging Approach. Biological Psychiatry: Cognitive Neuroscience and Neuroimaging 8, 262–270 (2023).

49. W. Sun et al., The anterior cingulate cortex directly enhances auditory cortical responses in air-puffing-facilitated flight behavior. Cell Reports 38, 110506 (2022).

50. E. Likhtik, J. M. Stujenske, M. A Topiwala, A. Z. Harris, J. A. Gordon, Prefrontal entrainment of amygdala activity signals safety in learned fear and innate anxiety. Nature Neuroscience 17, 106–113 (2014).

51. A. Burgos-Robles et al., Amygdala inputs to prefrontal cortex guide behavior amid conflicting cues of reward and punishment. Nature Neuroscience 20, 824–835 (2017).

52. J. E. LeDoux, R. Brown, A higher-order theory of emotional consciousness. Proceedings of the National Academy of Sciences 114, E2016–E2025 (2017).

53. T. Brosch, D. Sander, Comment: The Appraising Brain: Towards a Neuro-Cognitive Model of Appraisal Processes in Emotion. Emotion Review 5, 163-168 (2013).

54. H. Saarimäki et al., Discrete Neural Signatures of Basic Emotions. Cerebral Cortex 26, 2563–2573 (2015).

55. P. A. Kragel, M. C. Reddan, K. S. LaBar, T. D. Wager, Emotion schemas are embedded in the human visual system. Science Advances 5, eaaw4358 (2019).

56. G. Tononi, Consciousness as Integrated Information: a Provisional Manifesto. The Biological Bulletin 215, 216–242 (2008).

57. R. Brown, H. Lau, J. E. LeDoux, Understanding the Higher-Order Approach to Consciousness. Trends in Cognitive Sciences 23, 754–768 (2019).

58. B. J. Baars, “Global workspace theory of consciousness: toward a cognitive neuroscience of human experience” in Progress in Brain Research, S. Laureys, Ed. (Elsevier, 2005), vol. 150, pp. 45–53.

59. D. Mobbs, C. C. Hagan, T. Dalgleish, B. Silston, C. Prévost, The ecology of human fear: survival optimization and the nervous system. Frontiers in Neuroscience 9 (2015).

60. J. S. Feinstein et al., Fear and panic in humans with bilateral amygdala damage. Nature Neuroscience 16, 270–272 (2013).

61. Y. Tian, D. S. Margulies, M. Breakspear, A. Zalesky, Topographic organization of the human subcortex unveiled with functional connectivity gradients. Nature Neuroscience 23, 1421–1432 (2020).

62. W. M. Pauli, A. N. Nili, J. M. Tyszka, A high-resolution probabilistic in vivo atlas of human subcortical brain nuclei. Scientific Data 5, 180063 (2018).

63. D. Scheinost et al., Ten simple rules for predictive modeling of individual differences in neuroimaging. NeuroImage 193, 35–45 (2019).

64. R. A. Poldrack, G. Huckins, G. Varoquaux, Establishment of Best Practices for Evidence for Prediction: A Review. JAMA Psychiatry 77, 534–540 (2020).

65. B. T. T. Yeo et al., The organization of the human cerebral cortex estimated by intrinsic functional connectivity. Journal of Neurophysiology 106, 1125–1165 (2011).

## Supplementary References

1. V. Taschereau-Dumouchel, M. Kawato, H. Lau, Multivoxel pattern analysis reveals dissociations between subjective fear and its physiological correlates. Molecular Psychiatry 25, 2342–2354 (2020).

2. O. Esteban et al., fMRIPrep: a robust preprocessing pipeline for functional MRI. Nature Methods 16, 111–116 (2019).

3. K. Gorgolewski et al., Nipype: A Flexible, Lightweight and Extensible Neuroimaging Data Processing Framework in Python. Frontiers in Neuroinformatics 5 (2011).

4. B. B. Avants, N. Tustison, G. Song, Advanced normalization tools (ANTS). Insight j 2, 1–35 (2009).

5. Y. Zhang, M. Brady, S. Smith, Segmentation of brain MR images through a hidden Markov random field model and the expectation-maximization algorithm. IEEE Transactions on Medical Imaging 20, 45–57 (2001).

6. R. W. Cox, AFNI: Software for Analysis and Visualization of Functional Magnetic Resonance Neuroimages. Computers and Biomedical Research 29, 162–173 (1996).

7. D. N. Greve, B. Fischl, Accurate and robust brain image alignment using boundary-based registration. NeuroImage 48, 63–72 (2009).

8. F. Zhou et al., A distributed fMRI-based signature for the subjective experience of fear. Nature Communications 12, 6643 (2021).

9. L. J. Chang, P. J. Gianaros, S. B. Manuck, A. Krishnan, T. D. Wager, A Sensitive and Specific Neural Signature for Picture-Induced Negative Affect. PLOS Biology 13, e1002180 (2015).

10. M. Čeko, P. A. Kragel, C.-W. Woo, M. López-Solà, T. D. Wager, Common and stimulus-type-specific brain representations of negative affect. Nature Neuroscience 25, 760–770 (2022).

11. L. J. Chang et al., Endogenous variation in ventromedial prefrontal cortex state dynamics during naturalistic viewing reflects affective experience. Science Advances 7, eabf7129 (2021).

12. J. D. Power, K. A. Barnes, A. Z. Snyder, B. L. Schlaggar, S. E. Petersen, Spurious but systematic correlations in functional connectivity MRI networks arise from subject motion. NeuroImage 59, 2142–2154 (2012).

