## supplements for "Capturing dynamic fear experiences in naturalistic contexts: An ecologically valid fMRI signature integrating brain activation and connectivity"

**The VIFS and GNAS, but not the AFSS or PINES, predict video clip-induced short-term stable fear experience**

We found that the VIFS (r = 0.26, P = $2\times10^{-9}$) and GNAS (r = 0.15, P = $5\times10^{-4}$), but not AFSS (r = -0.06, P = 0.215) or PINES (r = 0.02, P = 0.706), significantly predicted movie clip-induced fear (Supplementary Fig. 1).

Given that the CAFE was developed on the training data in study 1, we additionally applied these signatures to the hold-out test data in study 1 to facilitate a fair comparison. We found similar prediction performances (VIFS: r = 0.35, P = $5\times10^{-11}$; GNAS: r = 0.18, P = 0.001; AFSS: r = -0.02, P = 0.676; PINES: r = 0.02, P = 0.774) as compared to the whole sample. Importantly, these prediction performances were substantially lower than that of the CAFE (r = 0.67; all permutation P of the differences were less than $1 \times 10^{-4}$).

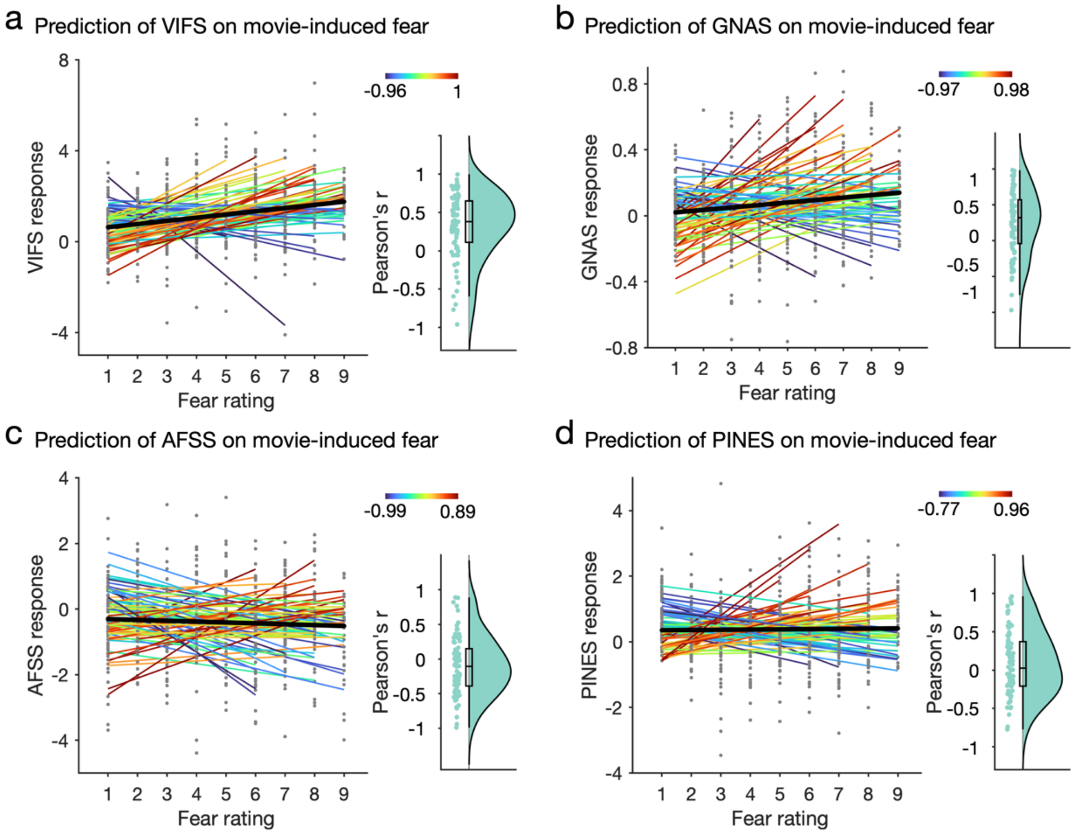


**Supplementary Fig. 1. Predicting movie clip-watching induced subjective fear experiences (study 1) with previously established activation-based signatures.** The VIFS (panel a; r = 0.26, P = $2\times10^{-9}$) and GNAS (panel b; r = 0.15, P = $5\times10^{-4}$) significantly predict movie clip-induced subjective feelings of fear whereas the AFSS (panel c; r = -0.06, P = 0.215)) and PINES (panel d; r = 0.02, P = 0.706) fail to predict movie-included subjective fear experience. VIFS, visually induced fear signature; GNAS, Generalized negative affect signature; AFSS, animal fear schema signature; PINES, picture-induced negative emotion signature.

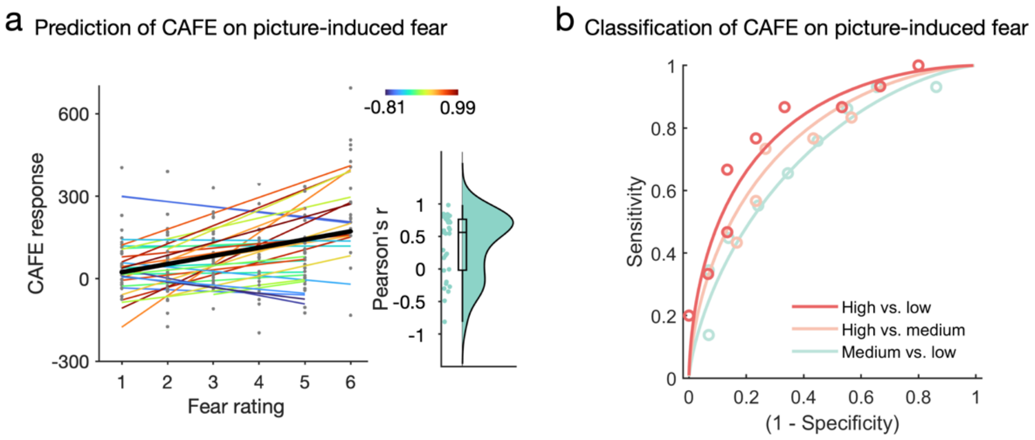


**Supplementary Fig. 2. Predicting picture-induced subjective fear experiences with the activation pattern of the CAFE.** a, the activation pattern of the CAFE accurately predicts subjective fear experience induced by animal pictures (overall prediction-outcome r = 0.35, P = $6\times10^{-6}$). Each colored line represents prediction within each individual participant. The black line indicates the overall (i.e., within and between participants) prediction. Raincloud plots show the distribution of within-participant predictions. b, the activation pattern of the CAFE can classify high fear (average of rating 4 and 5) from medium fear (average of rating 2 and 3; accuracy = 73%, P = 0.016, d = 0.74) and low fear (average of rating 0 and 1; accuracy = 77%, P = 0.005, d = 0.89). In addition, the CAFE to some extent distinguishes medium fear vs. low fear (d = 0.56) although the accuracy is not significantly higher than chance level (accuracy = 66%, P = 0.136).

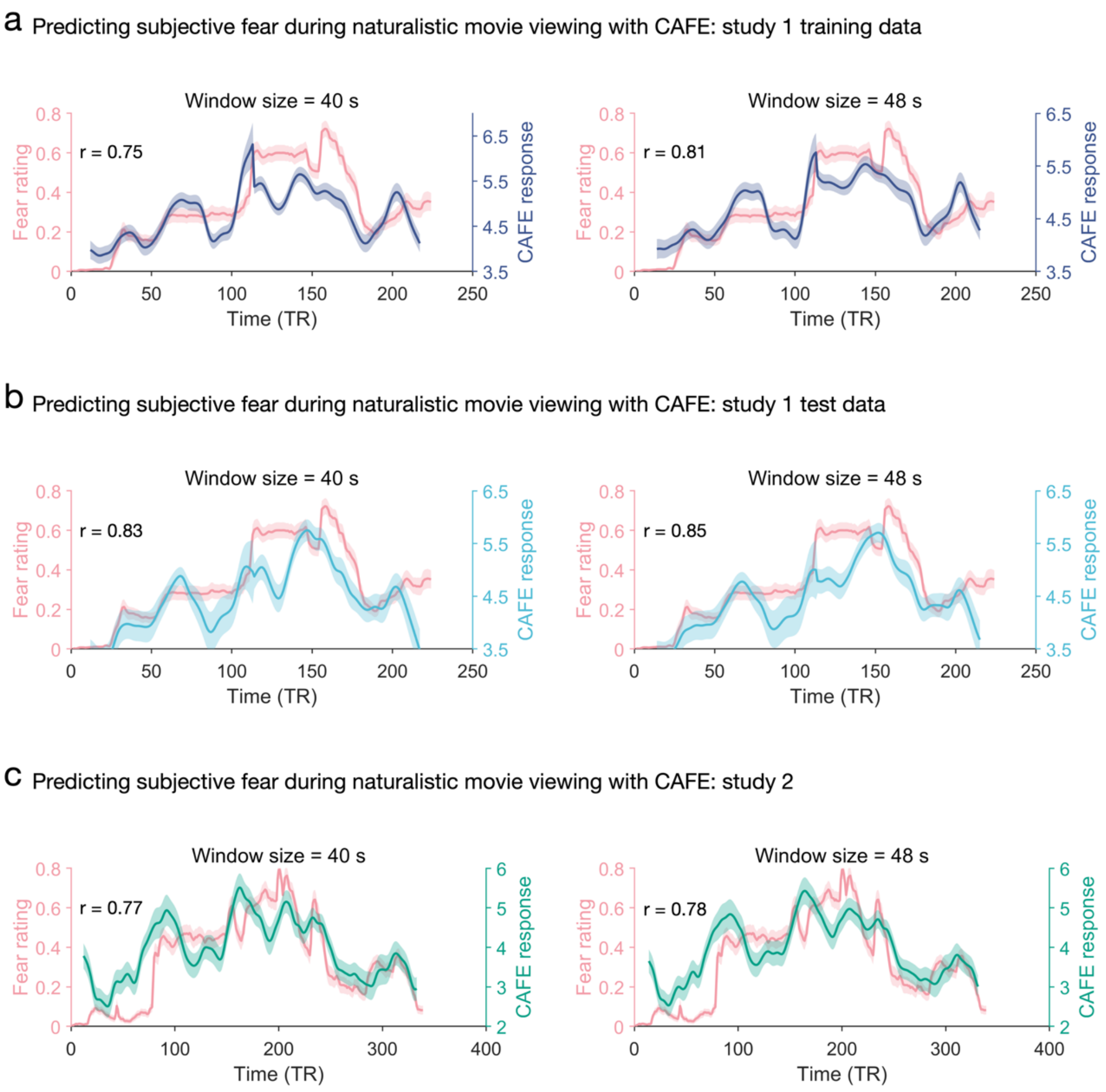


**Supplementary Fig. 3. Results of group-average subjective fear prediction from the CAFE using different window sizes.** The CAFE accurately tracks the dynamic fear changes using window sizes of 20 and 24 TRs in study 1 training dataset (a) and test dataset (b) as well as study 2 (c). These results suggest that our findings remain robust across selection of window sizes.

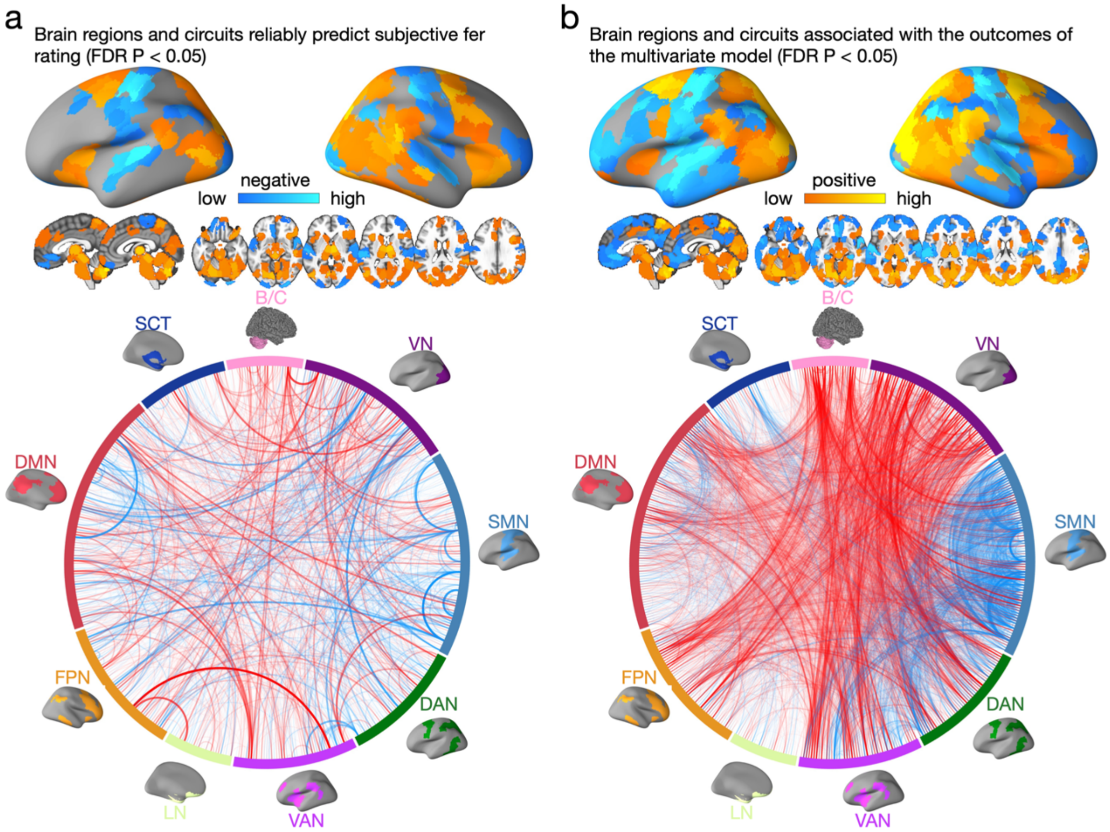


**Supplementary Fig. 4. Brain regions and pathways associated with and predictive of subjective fear rating.** a, model weight maps showing brain regions and circuits reliably predictive of fear ratings (FDR P < 0.05). Circular plots represent the significant pathways. b, model encoding maps showing brain regions and circuits reliably associated with the outcomes of the multivariate model (FDR P < 0.05). Circular plots represent the significant pathways.

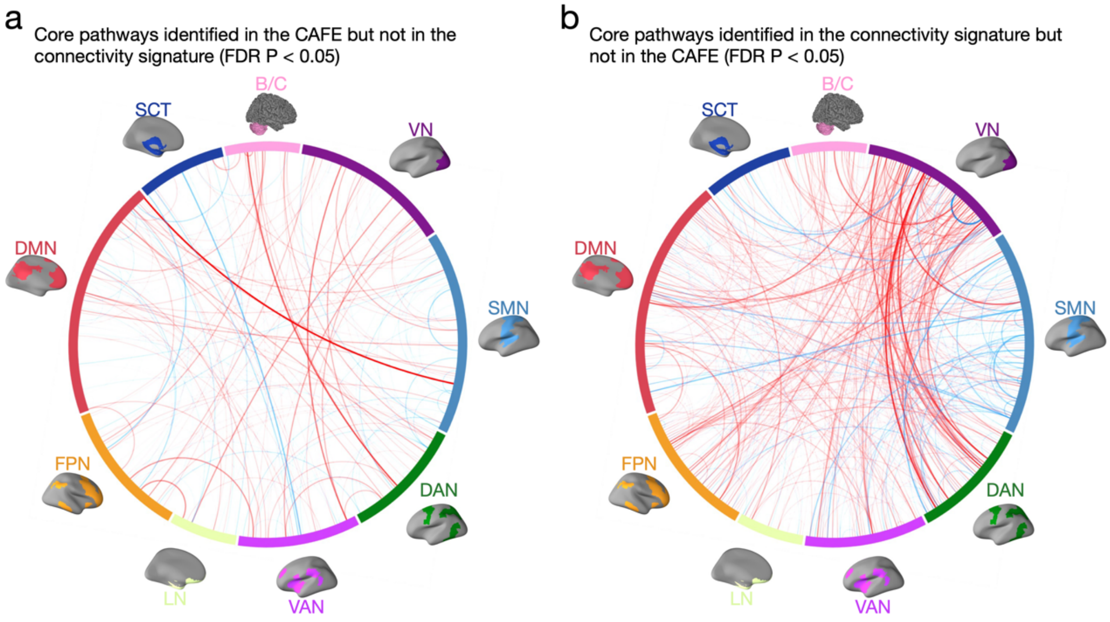


**Supplementary Fig. 5. Core brain pathways for the CAFE and the connectivity signature.** Circular plot visualizations of core pathways identified in the CAFE but not in the connectivity signature (panel a) and pathways identified in the connectivity signature but not in the CAFE (panel b); red line indicates positive weight and association and blue line indicates negative weight and association (all FDR $P < 0.05$).

**Supplementary Table 1 Predictive capacity of each large-scale brain network**

| $R^2_{total}$ | $R^2_{8net}$ | $R^2_{1net}$ | $R^2_{shared}$ | $\Delta R^2_{8net}$ | $\Delta R^2_{1net}$ | Single network |
|---|---|---|---|---|---|---|
| 0.421 | 0.403 | 0.291 | 0.273 | 0.130 | 0.018 | VN |
| 0.417 | 0.384 | 0.294 | 0.261 | 0.123 | 0.033 | SMN |
| 0.420 | 0.414 | 0.300 | 0.294 | 0.120 | 0.006 | DAN |
| 0.416 | 0.411 | 0.254 | 0.249 | 0.162 | 0.005 | VAN |
| 0.419 | 0.408 | 0.148 | 0.137 | 0.271 | 0.011 | LN |
| 0.416 | 0.410 | 0.240 | 0.234 | 0.176 | 0.006 | FPN |
| 0.429 | 0.418 | 0.288 | 0.277 | 0.141 | 0.011 | DMN |
| 0.419 | 0.409 | 0.168 | 0.157 | 0.251 | 0.010 | SCT |
| 0.419 | 0.409 | 0.254 | 0.245 | 0.164 | 0.009 | B/C |

Note. We performed a variance partitioning analysis to decompose the proportion of explained variance ($R^2_{total}$) to the variance explained only by the 8-network-model predictions ($\Delta R^2_{8net}$), variance explained only by the 1-network-model predictions ($\Delta R^2_{1net}$), and variance explained by predictions from both model trained on both 8 networks and model trained on the remaining 1 network ($R^2_{shared}$).
